# Generative AI Enables Breast Cancer Genomic Subtype Prediction from Histology Images

**DOI:** 10.64898/2025.12.29.692457

**Authors:** Brennan G. Simon, Clemens L. Weiss, Darren Chan, Lise Mangiante, Zhicheng Ma, Nicholas H. Smith, Allison Meisner, James M. Rae, Corey W. Speers, Kathy S. Albain, Cansu Karakas, Gregory R. Bean, Silvana Mouron, Miguel Quintela-Fandino, Christina Curtis

## Abstract

Breast cancer subtyping is essential for precision oncology, influencing prognosis, treatment selection, and clinical trial design. The Integrative Subtype Classification (IC) categorizes breast tumors into groups with distinct long-term outcomes based on genomic and correlated transcriptomic features. This method relies on sequencing data, which, despite decreasing costs, is not always available in research or clinical settings. Here we introduce PATH-IC, a computational pathology model that predicts ER+ breast cancer IC subtype risk of relapse categories from routine histology data. We enhance the current state-of-the-art computational pathology approach with BERGERON, which leverages generative AI to correct class imbalance and reduce overfitting, showing that synthetic data improves PATH-IC’s performance by the equivalent of 41% more real training samples. PATH-IC achieves a testing AUROC of 0.814, with predictions correlating to Oncotype DX scores and long-term relapse risk. Using attention-based model interpretation and CRAWFORD, a novel embedding-to-image foundation model, we demonstrate that PATH-IC identifies expected tumor microenvironment patterns for IC subtypes and highlights heterochromatin condensation as a key feature of high-risk tumors. Matched single-cell spatial transcriptomics confirm IC subtype-specific gene expression patterns identified by PATH-IC, including active metabolic, proliferative, and proteostasis pathways in the high-risk group. PATH-IC advances computational pathology through generative AI, enabling subtype inference from histopathology data.

## Introduction

The Integrative Subtype Classification (IC) system stratifies breast cancers into 11 distinct subtypes, characterized by distinct long-term relapse trajectories based on genomic alterations^1–3^. Among the approximately 80% of breast cancers classified as ER+, four subtypes are associated with a high risk of long-term (up to 20 year) relapse (IC1, IC2, IC6, and IC9), while four subtypes have typical relapse risks (IC3, IC4ER+, IC7, and IC8)^1,4^. The IC subtypes not only predict long-term relapse risk, but they also identify potentially targetable oncogene amplifications to inform treatment stratification. However, tumor sequencing data is currently required to determine IC subtype which is not always available in the research or clinical setting, especially in low-resource environments. Here, we introduce PATH-IC, a computational pathology model designed to predict High versus Typical Risk IC subgroups in ER+ breast cancers from routine histology images, enabling rapid and low-cost determination of this biomarker (Figure 1a).

**Figure 1.**
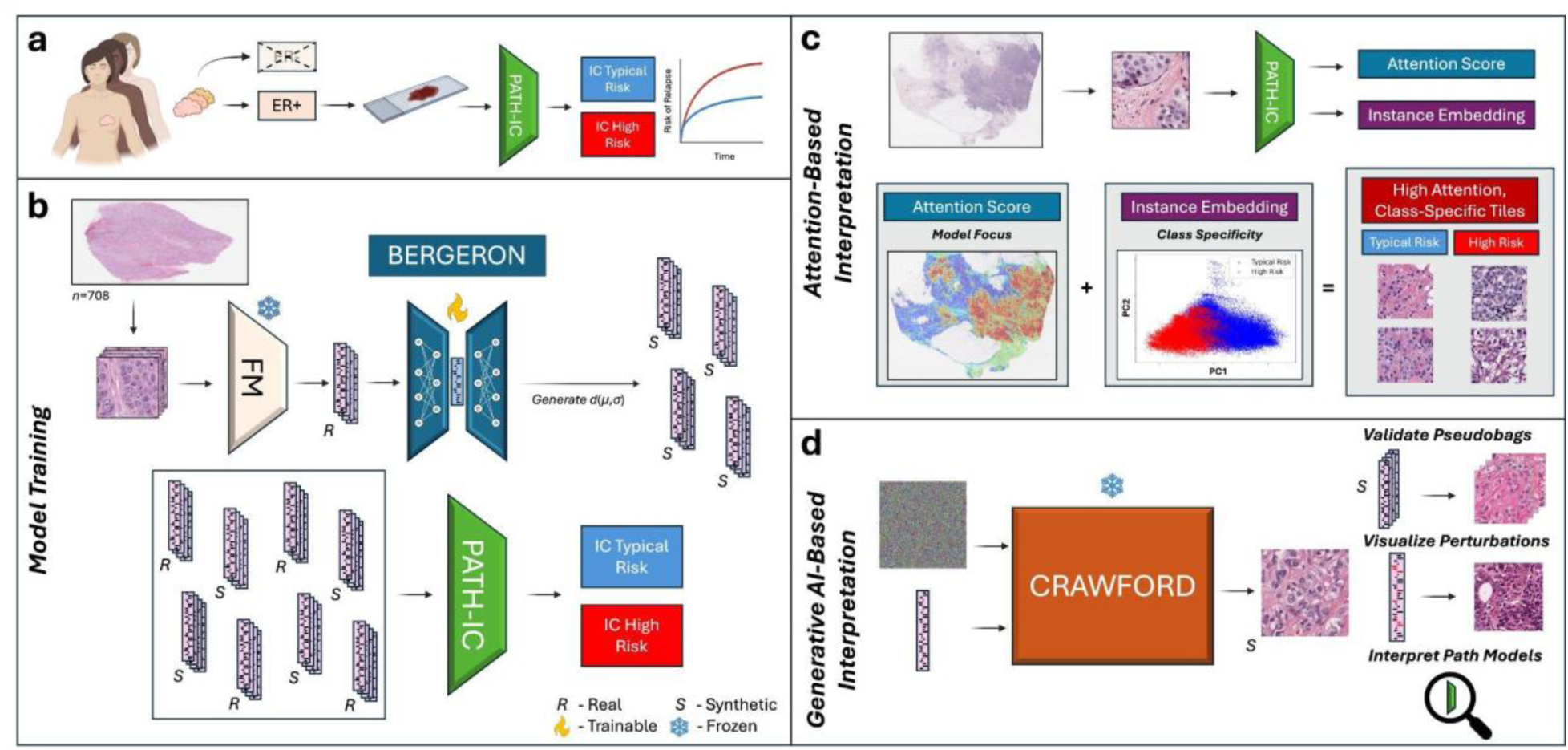
Schematic Overview of PATH-IC, BERGERON, and CRAWFORD. **a)** PATH-IC is a computational pathology model that predicts High versus Typical Risk of relapse Integrative Cluster (IC) subtypes amongst ER+ breast cancers using clinical histology data. **b)** PATH-IC was trained using the current state-of-the-art computational pipeline augmented by BERGERON, a generative AI framework we developed to overcome challenges of class imbalance and improve computational pathology model performance. Each training whole slide image was split into tiles and embeddings were extracted from the tiles using a pretrained foundation model (UNI2). BERGERON generates novel class-specific synthetic tile embeddings to increase training dataset size and balance classes. We combined BERGERON-produced synthetic tile embeddings with real tile embeddings to train PATH-IC, an attention-based multiple instance learning model (CLAM architecture), to predict ER+ IC subtype. **c)** Attention-based model interpretation traditionally involves visualizing the tiles to which the model assigns the highest attention scores. Attention scores represent the weight of each tile’s contribution to the overall slide class prediction, and high attention scores are interpretable as the tiles which the model focuses the most on in its prediction. Information from the instance embeddings of each tile can also be used to identify the tiles which the model finds the most class-specific, and when combined with attention scores, this approach can identify tiles that the model focuses the most on and finds the most class-specific. **d)** CRAWFORD, a foundation model for UNI2 embedding visualization, enables synthetic data validation and model interpretation. CRAWFORD is a conditional diffusion model which takes in as input a UNI2 tile embedding and a random noise image, and from these CRAWFORD produces a high-quality histology image that represents the tile embedding. We used CRAWFORD to validate the synthetic data we produced with BERGERON, visualize tile embedding perturbations, and interpret the knowledge base of PATH-IC.

Computational pathology is rapidly advancing due to developments in deep learning and computational hardware. Foundation models, such as UNI2, excel at converting histology images to vectorized embeddings, facilitating the design of diverse prediction models tailored for clinically relevant applications, including the inference of microsatellite instability (MSI) in colorectal cancer and predictions of mutation types and transcriptomic profiles^5–8^. Nonetheless, many computational pathology models face performance constraints owing to limited availability of training histological data^9,10^. For models reliant on matched patient data—such as genomic sequencing, RNA expression, or longitudinal outcome data—insufficient sample sizes can be a barrier to effective training^11^. Additionally, computational pathology models, akin to many deep learning applications, are susceptible to overfitting, particularly when trained on class-imbalanced datasets or smaller datasets^12,13^. The application of synthetic data in other domains of machine learning has proven effective in addressing these challenges by enhancing model generalization^14–16^. Concurrently, generative AI-based has been explored to alleviate data scarcity in computational pathology^17–19^, typically categorized into image-generating strategies that synthesize histology tile images and embedding-generating approaches that produce synthetic tile embeddings. While image-generating techniques can augment sample diversity, they are computationally intensive and prone to introducing artifacts. In contrast, embedding-generating methods offer greater efficiency but currently hinge on predefined augmentation strategies.

To mitigate data scarcity in the training of PATH-IC, we developed BERGERON, a class-conditional variational autoencoder that generates foundation model tile embeddings which are then assembled into novel synthetic histology samples (Figure 1b). By synthesizing tile embeddings and employing probabilistic latent space sampling, BERGERON allows for flexible data generation that mirrors the distributions of authentic samples and circumvents artifacts that arise in direct image generation. The seamless integration of BERGERON into established computational pathology workflows facilitates its application across a range of computational tasks.

Model interpretation is a critical aspect of computational pathology as it verifies that models recognize significant, non-spurious signals and facilitates biological discovery (Figure 1c)^20,21^. Accordingly, we have implemented multiple interpretive techniques for PATH-IC, including CRAWFORD, a UNI2 embedding-to-image foundation model. CRAWFORD enables the visualization of any UNI2 embedding, aiding the histological interpretation of differences in tile embeddings (Figure 1d). Our interpretive analysis indicates that PATH-IC discerns meaningful IC subtype-related signals, encompassing both known associations, such as microenvironment composition, as well as novel biological insights such as differences in nuclear morphology and differential enrichment of cellular energy and intracellular signaling pathways between subtypes. Through the application of BERGERON and CRAWFORD, we demonstrate that generative AI significantly enhances both the predictive performance and interpretation of breast cancer subtype classification in computational pathology, with the potential for broader applicability to other tasks with limited or imbalanced datasets.

## Results

### BERGERON Generates High Quality Synthetic Data

We initially trained a computational pathology model to predict ER+ breast cancer IC subtype using the UNI2 foundation model and CLAM, a state-of-the-art attention-based multiple instance learning (ABMIL) method for computational pathology, on data from the TCGA-BRCA cohort (*n*=708)^8,22,23^. CLAM training resulted in substantial overfitting in the High Risk IC subtype, indicated by a large difference between training and validation accuracy (Figure 2a). This overfitting is likely due to the large class imbalance between the High and Typical Risk groups and the rarity of High Risk IC subtype histology samples available for training (*n*=215, Figure 2b). Scarcity of training data with linked histology and genomics limits certain computational pathology applications, especially those involving rare subtypes. To address this, we developed BERGERON, a generative AI approach to produce additional labeled histology data to enrich our training dataset. BERGERON leverages a class-conditional variational autoencoder to produce high quality synthetic histology tile embeddings, as indicated by the similar distribution of real and synthetic data in principal component (PC) space (Figure 2c-d). Furthermore, BERGERON faithfully grouped synthetic tile embeddings to create realistic pseudobags, or collections of individual synthetic tiles that resemble a complete whole slide image (WSI), aided by its ability to consistently cluster tiles from the same WSI in the BERGERON latent space (Figure 2e-f, S1a-d).

**Figure 2.**
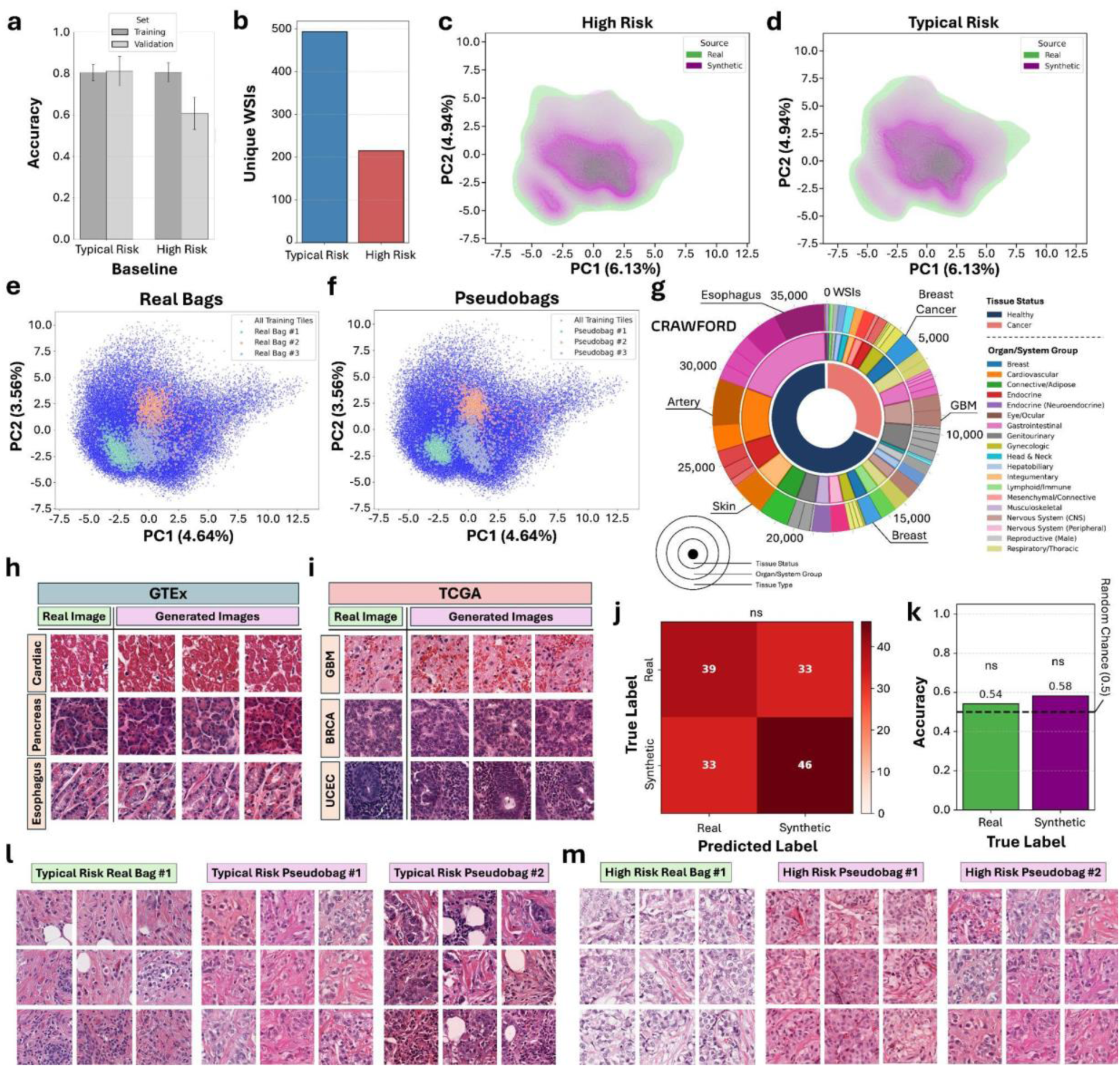
BERGERON and CRAWFORD Generate High Quality Synthetic Data. **a)** CLAM training and validation accuracy for Typical Risk and High Risk samples using real data. **b)** Unique Typical Risk and High Risk sample count used for training and validation. **c-d)** PCA coestimated on real tile embeddings (green) and BERGERON-produced synthetic tile embeddings (purple) for **c)** High Risk tiles and **d)** Typical Risk tiles. **e)** Example real bags, or the collections of tile embeddings of real WSIs. **f)** Corresponding pseudobags, or the collections of BERGERON-generated synthetic tile embeddings. **g)** Unique WSI counts used in training CRAWFORD. 100 tiles per WSI were used. **h-i)** Validation histology images and corresponding generated images by CRAWFORD for the GTEx and TCGA datasets. **j)** Breast pathologist confusion matrix for discriminating real and synthetic histology tiles. **k)** Pathologist accuracy for identifying real and synthetic histology tiles. **l-m)** Tiles from real WSI samples and tiles from generated pseudobags shown for **l)** Typical Risk and **m)** High Risk.

In order to visualize the synthetic tiles produced by BERGERON to validate that they resemble real breast cancer data, we developed CRAWFORD, a computational pathology foundation model that generates accurate histology tile images from UNI2 tile embeddings. Since UNI2 was trained on a wide variety of both healthy and malignant tissue, we similarly trained CRAWFORD on a comprehensive set of 61 healthy and malignant tissue types from the GTEx and TCGA datasets^23,24^. Altogether this included over 3.6 million tiles from 36,651 unique WSIs including 894 healthy breast samples and 1,105 breast cancer samples (Figure 2g, S2a). CRAWFORD generates highly accurate images based solely on input UNI2 embeddings across tissue types, with expected hue differences since UNI2 was trained to remove staining variance (Figure 2h-i). Given an even mixture of real and CRAWFORD-generated tiles across tissue types, an expert breast pathologist was unable to discriminate between the real and synthetic tiles, validating CRAWFORD’s performance (Figure 2j-k). We then used CRAWFORD to visualize pseudobags produced by BERGERON and found that the pseudobags both mirror Typical Risk and High Risk breast cancer histology and share consistent staining and morphologic patterns (Figure 2l-m). These data indicate that BERGERON and CRAWFORD generate high quality synthetic histology data.

### BERGERON Improves PATH-IC’s Performance

We next added the BERGERON-produced pseudobags to the existing real training data and retrained the CLAM model on this enriched dataset, resulting in a model termed PATH-IC. Increasing the amount of synthetic data in the training dataset led to a consistent improvement in model performance (Figure S3a-b), but exceeding a 10-fold increase resulted in a slight decline, indicating a trade-off between the benefits of synthetic data and the diminishing impact of real data. Optimal model performance used the full real training dataset and 5,400 pseudobags, 2,700 from each class with each bag containing 1,000 synthetic tile embeddings. This approach yielded an average cross-fold validation AUROC of 0.795 which, compared to the baseline CLAM model (trained without synthetic data), yielded notable performance improvements (Figure 3a, S4a). Compared to the baseline model, PATH-IC had an 8.9% improvement in High Risk validation accuracy and a 1.0% improvement in AUROC (Figure 3b,d, S4b). Typical Risk validation accuracy fell to 76.2%, which is expected given that the baseline model’s validation accuracy being higher than its training accuracy reflected model overfitting and inflated performance (Figure 3c). In contrast to the baseline model, PATH-IC displayed a healthy balance of High Risk and Typical Risk prediction performance indicated by significantly smaller gaps in three measures of model overfitting (Figure S4c-e). We also evaluated SEQUOIA, a recent model designed to predict bulk mRNA expression from histology, for its ability to predict IC subtypes, and we found that both PATH-IC and the baseline model significantly surpassed this non-ABMIL approach (Figure S3c-k). In order to evaluate the added value of BERGERON, we progressively subsampled the real WSIs used in training the baseline model and measured model performance, then fit a function of real training data to expected model AUROC for this task. We found that the 1.0% AUROC improvement using BERGERON is equivalent to the expected performance of training the model with an additional 292 real WSIs, or 41.2% of the entire TCGA-BRCA training dataset (*n*=708, Figure 3e).

**Figure 3.**
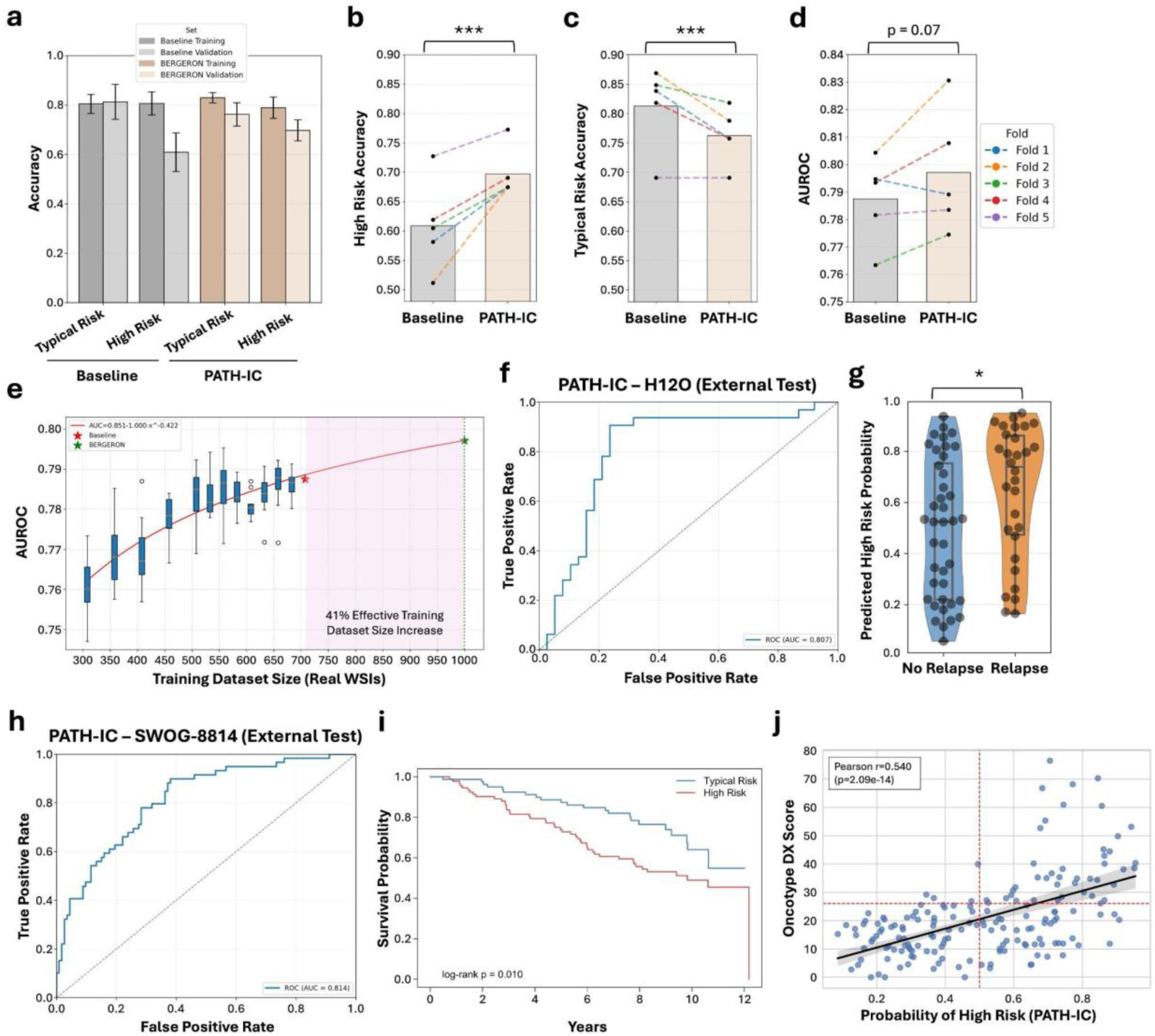
BERGERON Improves PATH-IC’s Performance. **a)** Class-specific training and validation accuracy for the baseline model (no BERGERON, real data only) and PATH-IC (using BERGERON-produced synthetic data). **b-d)** Five-fold cross validation comparison of baseline and PATH-IC in **b)** High Risk validation accuracy, **c)** Typical Risk validation accuracy, and **d)** validation AUROC. Connected data points indicate matched folds including identical real training and validation data with the addition of synthetic training data in the BERGERON condition. **e)** Model validation AUROC with variable amounts of real training data. Complete baseline model performance indicated with a red star and PATH-IC AUROC shown with a green star. **f)** PATH-IC testing ROC curve for the external H12O cohort (*n*=70). **g)** Predicted High Risk IC subtype probability for samples from patients with and without relapse in the H12O cohort. **h)** PATH-IC testing ROC curve for the external SWOG-8814 cohort (*n*=172). **i)** Kaplan-Meier curve showing disease-free survival by PATH-IC predicted IC subtype in the SWOG-8814 cohort. **j)** Scatterplot of PATH-IC High Risk prediction probability and inferred Oncotype DX score in the SWOG-8814 cohort.

We evaluated PATH-IC on two external cohorts with matched histology and ground truth IC subtype, the H12O cohort (*n*=70) and the SWOG-8814 clinical trial cohort (*n*=172). We found similar AUROC scores of 0.807 and 0.814 respectively (Figure 3f,h, S5a-i). PATH-IC’s performance in the SWOG-8814 cohort was better than the baseline model’s performance, as indicated by a higher AUROC, although both external cohorts were underpowered for this test as roughly 500 real WSIs would be needed to detect a 1.0% AUROC difference at 80% power (Figure S5j). Leveraging long-term follow-up data (median 11.4 years) in the H12O cohort, we found that patients who experienced long-term relapse were more likely to have a High Risk subtype prediction from PATH-IC, consistent with the IC subtypes classification (Figure 3g, S6a-b,d-e). A multivariate Cox proportional hazards model adjusted for clinical covariates indicates that PATH-IC predicted High Risk probability was associated with an elevated relapse hazard ratio in the H12O cohort (HR=7.50, 95% CI: 1.29-43.57, *p*=0.02) (Figure S6c). Patients in the SWOG-8814 cohort with High Risk PATH-IC predictions had significantly worse disease-free survival (HR: 1.926, 95% CI: 1.156, 3.210) (Figure 3i). PATH-IC predictions also correlated with inferred Oncotype DX scores in the SWOG-8814 cohort, revealing an enrichment of predicted High Risk samples with Oncotype DX scores above 26, consistent with the historical classification of scores greater than or equal to 26 being high-risk (Pearson r=0.540, *p*=2.09×10^-14^) (Figure 3j, S6f-g)^25,26^. Thus PATH-IC accurately predicts IC subtype risk categories and long-term breast cancer relapse risk.

### PATH-IC Model Interpretation Validates Known Features and Discovers New Signals

We next implemented model interpretation techniques to validate PATH-IC’s learning and derive new biological insight from the model. Standard model interpretation generally involves visualizing the tiles with the highest attention scores computed during ABMIL training. However, this approach did not illuminate class differences between the top 100 attended High and Typical Risk tiles from each correctly predicted WSI, indicated by a strong correlation in attention scores for both subtype classes (Figure 4a). We hypothesized that this strong correlation in class attention scores indicates that PATH-IC uses attention scores to upweight tiles with high cancer density and that the class-differentiable signal was present elsewhere, supported by a positive correlation between attention score and cancer cell density for both classes (Figure S7a-b). Instead, we turned to the instance embeddings, the task-specific tile embeddings computed during CLAM training (Figure S8a-b). The instance embeddings of tiles with high attention scores displayed a strong separation by IC subtype in PC space (Figure 4b). We then visualized the top 50 most extreme tiles along PC1 of the instance embedding PCA and interpreted these tiles as the most class-distinctive high-attention tiles (Figure 4c-d). There were clear pathologic differences between the tiles that PATH-IC considers strongly High Risk and Typical Risk, notably a high density of cancer cells with irregular nuclei in the High Risk tiles and the presence of stromal and immune cells in the Typical Risk tiles. Review of the top 200 class-distinctive high-attention tiles by an expert breast pathologist found a more aggressive phenotype as well as higher rates of mitosis and necrosis in the High Risk tiles (Figure 4e). To scale this analysis to more tiles, we next quantified the cell types present in the top 2,000 class-distinctive high-attention tiles using HoVer-Net^27^, a pretrained cell type prediction model, and found that High Risk tiles have 1.7 times the proportion of cancer cells per tile as Typical Risk tiles, while Typical Risk tiles have 5.7 times the proportion of immune cells and 4.7 times the proportion of stromal cells per tile (Figure 4f). This matches the expected tumor microenvironment composition found in our recent large-scale transcriptomics analysis across the IC subtypes^3^.

**Figure 4.**
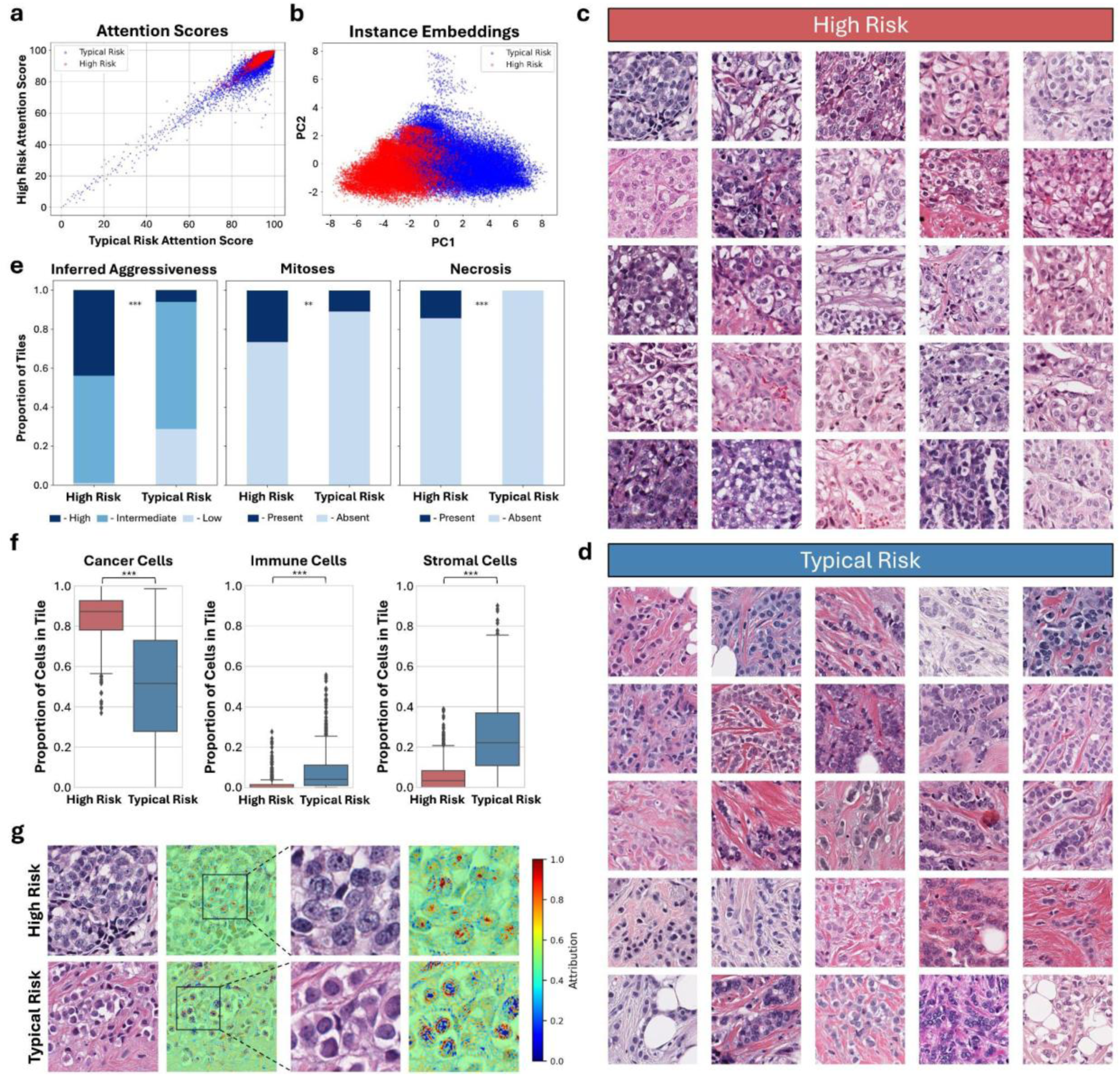
PATH-IC Attention-Based Model Interpretation. **a)** Typical Risk attention score and High Risk attention score for each of the top 100 attended tiles in correctly predicted samples colored by class. **b)** A PCA of each top attended tile’s instance embedding colored by class. **c-d)** The top 25 high attention score tiles with the highest and lowest instance embedding PC1 values, hereafter referred to as class-distinctive high-attention tiles for **c)** High Risk tumors and **d)** Typical Risk tumors. **e)** Pathologist-derived quantification of inferred aggressiveness, mitosis, and necrosis in the top 200 class-distinctive high-attention tiles from High and Typical Risk samples. **f)** Cell type proportions of the top 1,000 class-distinctive high-attention tiles for each subtype. Cell type predictions performed using HoVer-Net. **g)** Pixel level attention attribution scores for representative High Risk and Typical Risk class-distinctive high-attention tiles computed using Integrated Gradients.

Finally, we performed pixel-level attention analysis using Integrated Gradients^28^ to extend our tile-level attention analysis. We found that PATH-IC focused strongly on the nuclear interior, specifically prominent nucleoli and compact chromatin aggregates, when classifying tiles as High Risk and focused on the nuclear periphery when classifying tiles as Typical Risk, features not previously known to associate with IC subtype (Figure 4g). Attention analysis on the pseudobags generated by BERGERON revealed patterns consistent with the real data, including a high density of cancer cells in the High Risk tiles, the presence of immune and stromal cells in the Typical Risk tiles, and the same pixel-level attention patterns, indicating that BERGERON is capable of producing high quality synthetic data that CRAWFORD can accurately visualize (Figure S9a-c). Taken together, these results indicate that PATH-IC learned previously known and previously unknown histological features associated with IC risk groups.

### Perturbational Model Interpretation Reveals Smooth Feature Transitions

To identify the relative IC subtype specificity of histologic features identified in class-distinctive high-attention tiles, we next visualized the dynamic transition of these features from High Risk tiles to Typical Risk tiles across the instance embedding PC1 axis. Instead of tracking real tiles across the PC1 axis—which would require comparing tiles from different WSIs and thus introduce noise from staining variation, morphology differences, and other confounders—we developed a perturbational approach specifically designed to maximize the signal-driven histological changes as a tile moved from one extreme of the PC1 axis to the other. To do this, we took High or Typical Risk tile embeddings with extreme instance embedding PC1 values as input and applied gradient descent-based perturbations to modify the tile’s embedding until its PC1 value approached the opposite extreme (Figure 5a,d). Crucially, perturbations were only rewarded if they shifted the embedding in the desired PC1 direction, ensuring that each modification added signal and minimized noise. We then generated the associated images from the initial tile embedding and the perturbational class-switched embedding using CRAWFORD. This framework allowed us to evaluate how tiles would change if their embedding shifted toward the other class. This approach avoids the need to infer histological changes across tiles from different WSIs using real data and instead supports the isolation and visualization of class-defining features while holding all other sources of variability constant.

**Figure 5.**
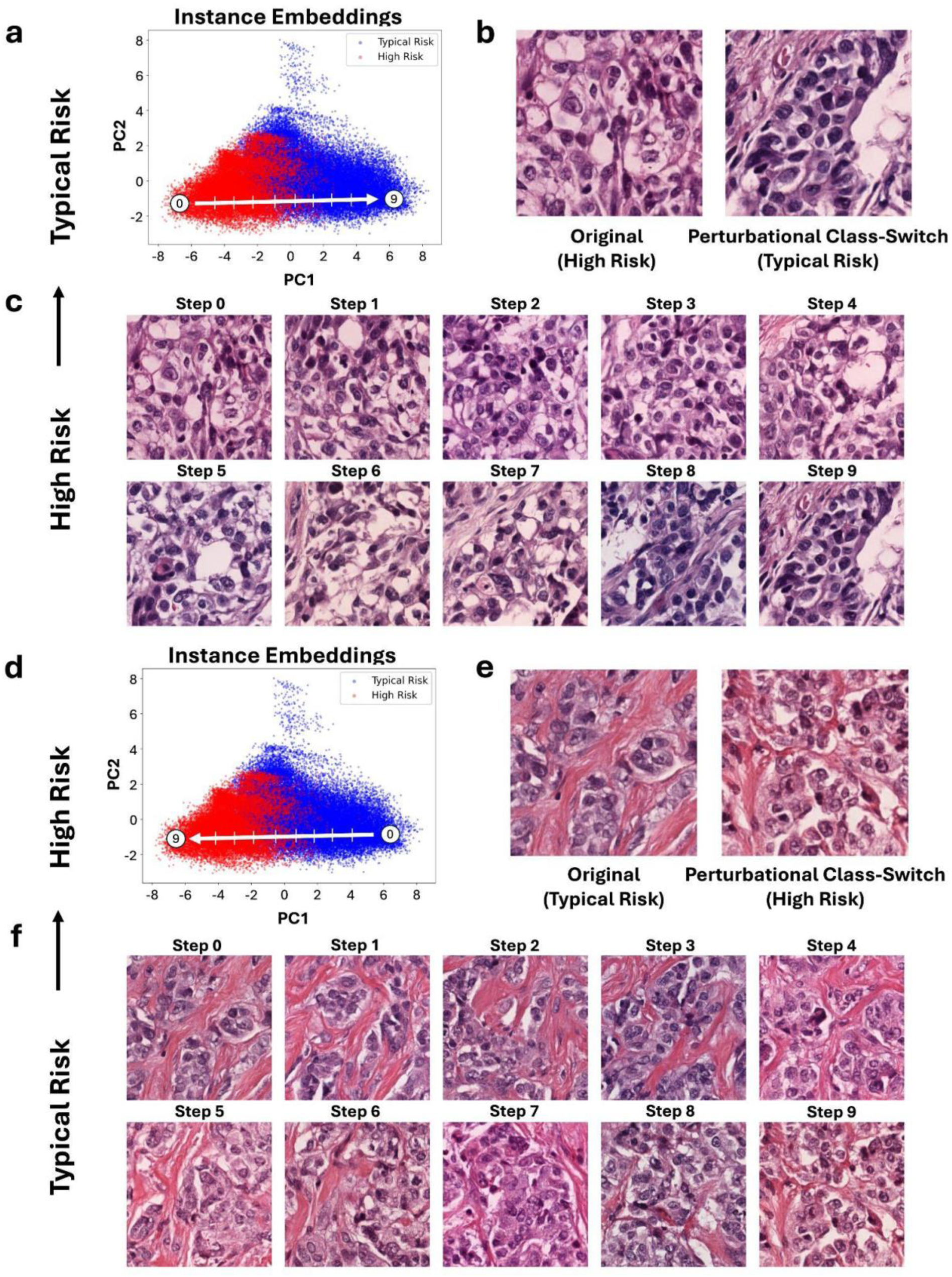
Perturbational Model Interpretation. **a)** PCA of the instance embedding of the top attended tiles indicating the selected High Risk start tile, Typical Risk end point, and direction of perturbation. **b)** Initial High Risk and perturbational class-switched tiles. Images generated using CRAWFORD. **c)** The full perturbational walk from the initial High Risk tile to the perturbational class-switched tile. Images generated using CRAWFORD. **d)** PCA of the instance embedding of the top attended tiles indicating the selected Typical Risk start tile, High Risk end point, and direction of perturbation. **e)** Initial Typical Risk and perturbational class-switched tiles. Images generated using CRAWFORD. **f)** The full perturbational walk from the initial Typical Risk tile to the perturbational class-switched tile. Images generated using CRAWFORD.

In class-switching from High Risk to Typical Risk, the initial tiles became more organized with cancer cells surrounded by an intact tumor-stroma boundary and stromal cells, indicating a lower grade phenotype (Figure 5b). Linear interpolation between these instance embeddings tracked a smooth path between High Risk features and Typical Risk features which we visualized using CRAWFORD (Figure 5c). The segmentation of cancerous regions from stromal regions happened relatively early in the transition of High Risk to Typical Risk, suggesting that the loss of stromal organization around the cancer cells is an indicator of high-confidence High Risk regions. Conversely, in class-switching from Typical Risk to High Risk, the number of cancer cells in the segmented tumor nodules of the initial tiles quickly grew indicating that increased density of cancer cells within their nodules is a more common feature to both Typical and High Risk tiles, whereas the speckling of the nuclei through chromatin condensation, which happened in later steps, is indicative of high confidence High Risk tiles (Figure 5e-f).

### Attention-Based Single Cell Spatial Transcriptomics Analysis Identifies IC Subtype-Specific Gene Expression Differences

To identify cancer cell-specific gene expression differences between High and Typical Risk IC subtype tumors, we turned to an internal atlas of matched single cell spatial transcriptomics (scST) and histology samples, consisting of coregistered biopsy cores from 204 patients from the H12O cohort assembled onto 9 tissue microarrays (TMA). We first deployed PATH-IC on the digitized histology slides and identified the tiles with the 1,000 most positive and 1,000 most negative instance embedding PC1 values (Figure 6a-b). We extracted all scST cells from the same regions as the class-distinctive tiles, filtered out non-cancer cells, and performed batch correction to normalize the data across tumors (Figure 6c-d). This resulted in a collection of cancer cells highly representative of the High Risk IC subtype and a separate collection of representative Typical Risk cancer cells. We then performed differential gene expression analysis at the pseudobulk level between the two groups and found a strong enrichment of genes that are part of the ground truth IC subtype classifier, iC10^1,29^, validating that PATH-IC learned true IC subtype biology (Figure 6e). We also found differentially expressed genes not previously known to be strongly associated with IC subtype, including *OXR1, COX6C*, and *NDUFB9* in the High Risk cancer cells and *CYP46A1, TLR3*, and *C3AR1* in the Typical Risk cancer cells.

**Figure 6.**
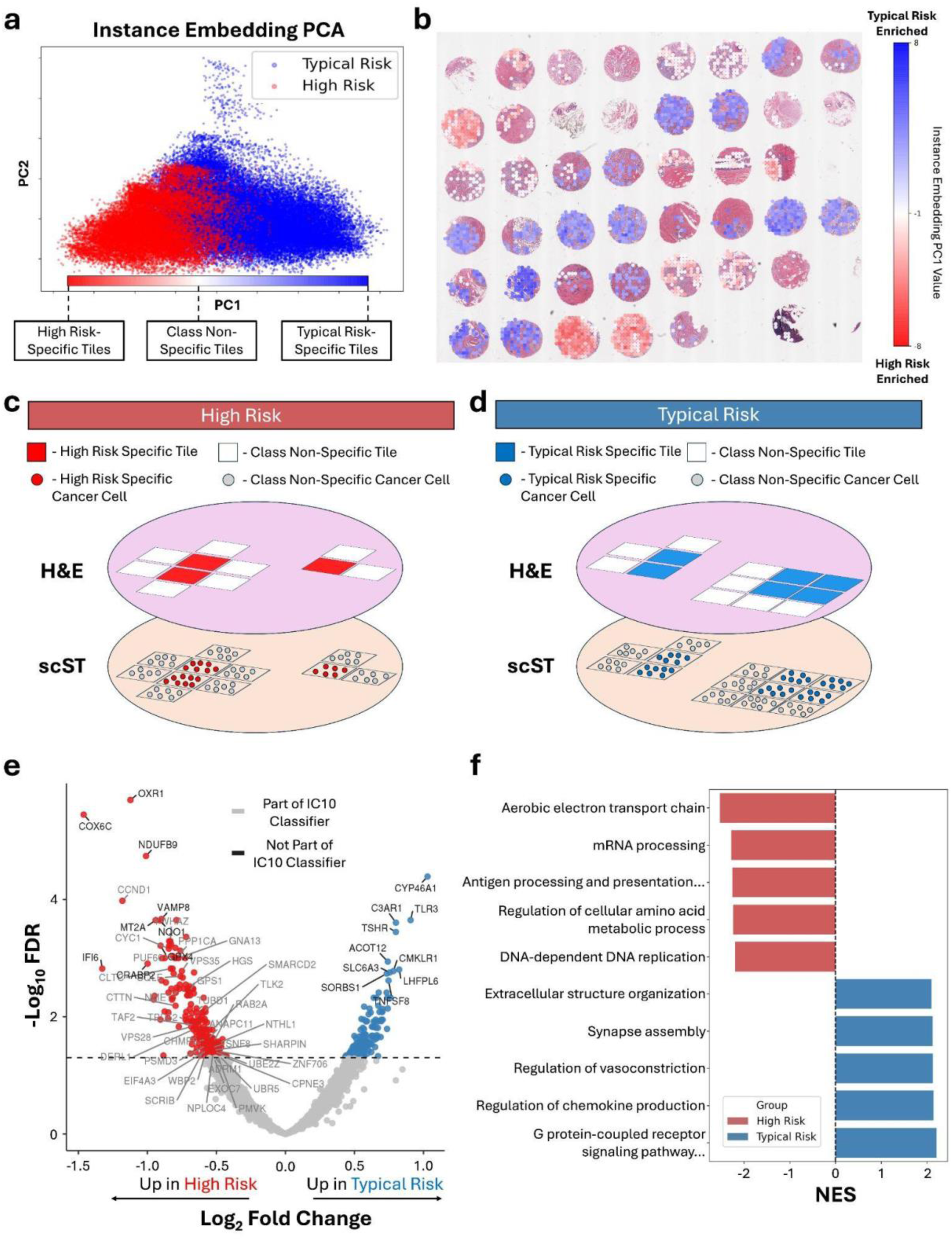
Attention-Based Single Cell Spatial Transcriptomics Analysis. **a)** An interpretation of the instance embedding PCA plot outlining PC1 scores and their relation to class specificity. **b)** Instance embedding PC1 values on a representative tissue microarray (TMA) from the H12O cohort. **c-d)** Schematics outlining coregistered histology (H&E) and single cell spatial transcriptomics (scST) samples and the selection of **c)** High and **d)** Typical Risk specific cancer cells. Histology tiles with extreme instance embedding PC1 values are interpreted as highly class-specific. Cancer cells from the scST dataset in the same regions as class-specific histology tiles are collected for further analysis. **e)** Volcano plot identifying individual genes enriched in class-specific cancer cells of High Risk and Typical Risk samples. Genes labeled in gray are included in the ground truth IC (iC10) classifier. **f)** Gene set enrichment analysis indicating the normalized enrichment score (NES) of representative enriched pathways for High Risk and Typical Risk cancer cells.

We then performed gene set enrichment analysis and found strong class specific enrichment of pathways (Figure 6f, S10a-b). Gene sets involved in oxidative phosphorylation, RNA processing, DNA replication, and proteostasis were strongly enriched in the High Risk cancer cells, representing possible targetable axes of this subtype. Conversely, gene sets relevant to G protein-coupled receptor signaling pathways were enriched in Typical Risk cancer cells, suggesting a subtype-specific mode of predominant intracellular signaling. Typical Risk cancer cells also maintained higher expression of pathways relevant to extracellular matrix organization and immune activation, reflecting a greater interaction with the stromal and immune microenvironments. To validate these gene expression differences between subtypes and demonstrate the utility of this analysis in circumstances without matched scST data, we imputed bulk mRNA expression from the top 1,000 class-distinctive high-attention tiles for each subtype from the TCGA-BRCA training dataset using SEQUOIA^7^ and performed gene set enrichment analysis (Figure S11a-b). Enriched gene sets in High Risk tiles included cell replication pathways and RNA processing pathways, similarly to the gene sets found in scST analysis. Many of the enriched gene sets in the Typical Risk tiles were related to extracellular matrix assembly and stromal gene programs which reflects the greater stromal involvement in the Typical Risk tiles.

### Whole Slide Attention Analysis Reveals the Complexity of IC Subtypes

We next extended our interpretation to the WSI level by visualizing patterns of instance embedding values throughout entire tumors. We binned each tile by its instance embedding PC1 value and represented tumors by their proportions of tiles in each bin. We found five major tumor clusters: two clusters corresponding to high-confidence Typical Risk tumors (Clusters 1 and 2), one cluster of high-confidence High Risk tumors (Cluster 5), and two clusters of heterogeneous and low confidence tumors (Clusters 3 and 4) (Figure 7a). Clusters 3 and 4, which account for 33.8% of the cohort, had notably lower mean prediction probabilities, high levels of class-indiscriminate tiles (bin 3), and often contained moderate to high-confidence tiles for both High Risk and Typical Risk. Visualization of instance embedding scores overlaid on the WSIs confirmed the presence of high-confidence tumors composed of many tiles with high confidence for High Risk or Typical Risk (Figure 7b), heterogeneous tumors with a mix of tiles with high confidence High Risk and Typical Risk features (Figure 7c), and low confidence tumors with large quantities of class indiscriminate tiles (Figure 7d). Inaccurate predictions for heterogeneous tumors may reflect a greater underlying subtype heterogeneity than expected for these samples and their binary IC subtype label. This analysis revealed a spectrum of IC subtype heterogeneity across ER+ breast cancer samples that warrants further investigation.

**Figure 7.**
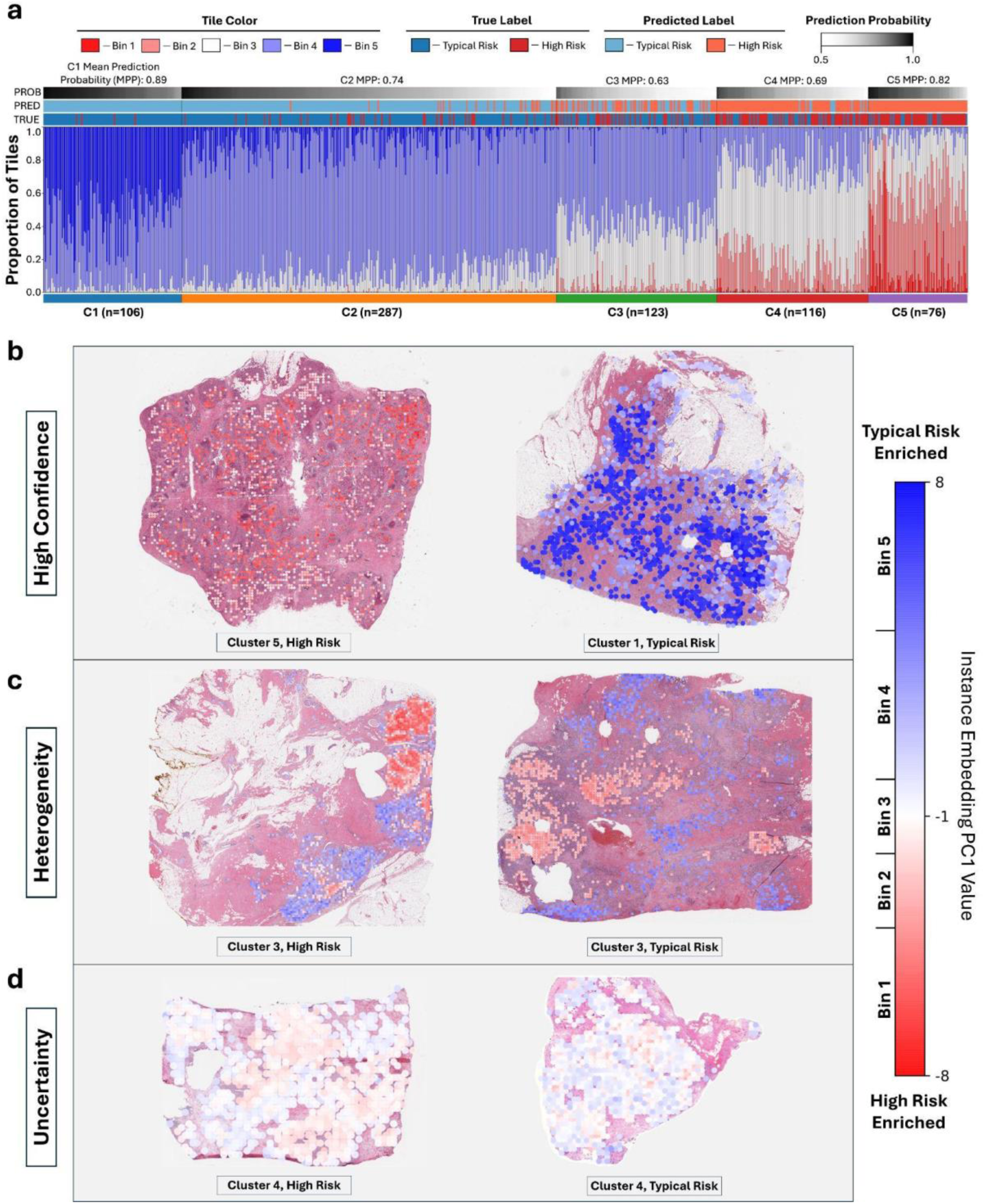
Whole Slide Attention Analysis. **a)** Waterfall plot depicting the proportion of each tumor’s tiles binned into groups based on tile instance embedding PC1 values. Unsupervised clustering was performed on the proportion of tiles in each bin for each tumor to determine tumor clusters. **b-d)** The 1,000 tiles with the highest instance embedding PC1 values and the 1,000 tiles with the lowest instance embedding PC1 values are plotted for each tumor. **b)** Example high-confidence tumors. **c)** Example heterogeneous tumors with a mix of High Risk tiles and Typical Risk tiles. **d)** Example tumors which lack class discriminative tiles.

## Discussion

Here we present PATH-IC, a computational pathology model that predicts IC subtype risk groups amongst ER+ breast cancer from histology images. Our findings indicate that PATH-IC’s predictions correlate with long-term patient relapse risk and Oncotype DX scores across two external cohorts. Notably, all samples with high-confidence Typical Risk predictions from PATH-IC (greater than 75% probability) had Oncotype DX scores below the standard threshold of 26^25,26^. This suggests that PATH-IC could serve as a rapid screening tool to triage cases requiring Oncotype DX testing.

We observed that the current state-of-the-art computational pathology approach led to significant overfitting in the High Risk subtype due to the limited training data available for this subgroup. This challenge is characteristic of many computational pathology tasks where one or more classes are underrepresented or imbalanced because of their rarity. Our results indicate that generative AI can effectively mitigate model overfitting for these underrepresented subtypes, in our case improving High Risk accuracy by 9%, which is comparable to the effect of adding 41% more real training samples. BERGERON is an attractive solution to the data scarcity problem prevalent in computational pathology, and by fitting conveniently into the standard computational pathology pipeline, BERGERON can be easily implemented to a wide range of tasks.

Our improved model interpretation ranks tiles by their attention as well as their embedding and leverages a new histology foundation model, CRAWFORD, to enable perturbational interpretation. Furthermore, we applied recent pretrained histology models HoVer-Net^27^ and SEQUOIA^7^ for cell-type prediction and gene expression imputation from histology, while Integrated Gradients^28^ enabled precise pixel-level model interpretation. Using these approaches, we found that PATH-IC learned expected biological features and discovered previously unknown histological signals associated with IC subtypes. The presence of more immune and stromal cells in class-distinctive high-attention Typical Risk tiles aligns with recent findings from bulk RNA sequencing analysis, which identified the Typical Risk microenvironment as biased toward the Immune-Fibrotic Enriched type, while the High Risk environment is enriched for the Depleted type^3,30^.

Analysis using Integrated Gradients further highlighted a pattern of model attention on nuclear speckles in High Risk tumors. These speckles often correspond to prominent nucleoli or heterochromatin aggregates. These heterochromatin aggregates may reflect broad genomic instability and ecDNA accumulation, features characteristic of High Risk tumors^3,31^. By leveraging PATH-IC’s attention-based learning alongside a dataset of matched histology and scST data, we uncovered subtype-specific gene expression differences in cancer cells. The enriched pathways suggest that High Risk tumors upregulate cellular energy, cell replication, and proteostasis pathways, consistent with increased rates of mitosis and cancer cell density in class-distinctive high-attention tiles. In situations where matched scST data are unavailable, recent methods have shown promise in predicting transcriptomic expression from histology, which when combined with attention-based learning, can identify gene expression patterns in class-specific tumor regions from histology images alone^7,32,33^.

Additionally, we identified five distinct patterns among ER+ breast tumors: three clearly corresponded to specific IC subgroups, while two exhibited mixed characteristics, indicating a higher proportion of heterogeneous samples than previously recognized. This spatial heterogeneity may contribute to PATH-IC’s inaccuracies, as the ground truth IC subtype was determined using bulk tissue sequencing, which, due to sample collection limitations, may have missed key areas of High Risk or Typical Risk signals that are present in the histology WSIs. These findings illuminate heterogeneity within tumors that warrant further investigation in relation to other molecular features and clinical outcomes and will be the focus of investigation in larger cohorts with linked WSI and omic data.

## Methods

### Data Preprocessing

PATH-IC was trained on the TCGA-BRCA dataset^23^. Patient samples were filtered by clinical ER positivity and multiple WSIs from the same patient were combined. WSI tiling was performed using the CLAM framework to create non-overlapping 128μm x 128μm tiles. Tiles were then filtered to remove poor quality tiles and tiles that contained no cancer cells. To do so, a ResNet50 model was trained to separate manually labeled high- and low-quality tiles and applied to the complete dataset. Patient samples with fewer than 400 remaining tiles (16) were removed, resulting in a total number of 708 unique patient samples for training. IC subtype was determined using EniCLust^3^. Sample labels for PATH-IC were determined by grouping EniCLust predictions into their respective categories: IC1, IC2, IC6, and IC9 were grouped into “High Risk” and IC3, IC4ER+, IC7, and IC8 were grouped into “Typical Risk”. UNI2 was used to extract tile embeddings according to the CLAM pipeline^8,22^. Data processing was similarly performed on the H12O and SWOG-8814 cohorts for external testing. IC subtype was determined using the iC10 package for both external cohorts due to the lack of matched normal tissue sequencing.

### PATH-IC

PATH-IC is a standard CLAM attention-based multiple instance learning model. To train the baseline model, a clam_mb model with size ‘big’ was used with dropout = 0.25, regularization = 1e-5, learning rate = 1e-5, and otherwise default parameters (Table S1). We found weighted sampling to produce the best performance when training on real data only, likely due to the class imbalance in real training samples. To train PATH-IC, an identical model architecture was used, and the learning rate was decreased to 1e-6 to accommodate the larger training dataset, however we removed weighted sampling as we found it slightly decreased model performance once large amounts of synthetic data were used. Five-fold cross validation was used across the entire TCGA-BRCA training dataset with samples split into folds based on their individual IC subtype. Both the baseline model and PATH-IC were trained until the validation dataset experienced minimal loss for each fold.

### BERGERON

BERGERON is a class-conditional variational autoencoder (VAE) designed to generate synthetic histology tile embeddings in a label-dependent manner. During training, each input consists of a real tile embedding concatenated with its one-hot-encoded class label. These inputs are passed through an encoder network, which projects the data into a lower-dimensional latent space regularized to follow a multivariate Gaussian distribution. The encoded representations are then passed through a decoder, which reconstructs the original embedding dimensionality—in our case matching that of a UNI2 tile embedding (1,536 features)—conditioned on the class label.

The total loss function consists of two components: (1) the Kullback-Leibler (KL) divergence loss, which encourages the latent space to approximate a unit Gaussian distribution, and (2) the reconstruction loss, which penalizes differences between the original and reconstructed embeddings. The KL divergence is weighted by a scalar parameter β, which is annealed linearly from a low starting value across training epochs to prevent posterior collapse and encourage stable training (Table S2).

To generate synthetic tile embeddings, we sample from the Gaussian latent space and decode these samples conditioned on the desired class. We found that real tiles from the same WSI clustered in BERGERON latent space, so to form realistic pseudobags we sampled tiles in distributions that match the shape of real bags (Figure S1). To construct pseudobags, BERGERON first encodes all tiles from a randomly selected real WSI, computes the mean and variance across the tile latent vectors, and then draws 1,000 samples from the resulting distribution. These sampled latent vectors are then decoded into synthetic tile embeddings and grouped to produce a pseudobag conditioned on the desired class. This process is repeated for every pseudobag construction cycle.

To avoid data leakage, a separate instance of BERGERON was trained independently for each training fold of the PATH-IC dataset, using only the training data within that fold. This ensures that no validation data from PATH-IC was seen by BERGERON during training. To prevent overfitting to WSIs with large tile counts, we limit training to a maximum of 2,000 randomly selected tiles per WSI per epoch.

### CRAWFORD

CRAWFORD is a foundation model for the visualization of UNI2 embeddings. CRAWFORD is a diffusion model that takes in as input a UNI2 embedding and a random noise image, and from these it produces a 256×256 pixel histology image representing the embedding. The architecture of CRAWFORD was inspired by RNA-CDM, a conditional cascaded diffusion model^17^. In order to match the breadth of tissue types used for training UNI2, we trained CRAWFORD on 36,651 WSIs from 61 healthy and malignant tissue types from the GTEx and TCGA datasets (Figure 2g, S2a)^23,24^. 100 randomly selected tiles from each WSI were extracted, passed through UNI2 to obtain their embeddings, then used to train CRAWFORD. CRAWFORD was trained until its loss stabilized and the produced images were satisfactory which was approximately 1 epoch. Validation images used in Figure 2 are from held-out WSIs.

### Attention-Based Model Interpretation

Attention-based interpretation of PATH-IC was performed on three levels. At the tile level, raw attention scores for each tile in each real bag were extracted using PATH-IC as well as each tile’s instance embedding. The top 100 tiles from each correctly predicted real sample with the highest correct class attention scores were accumulated, then from this collection the tiles with the most positive and most negative PC1 values from a PCA fit on the instance embedding vectors were considered class-distinctive high-attention tiles. Pathologist review of histological features was performed on the top 100 class-distinctive high-attention tiles from each class. The top 1,000 class-distinctive high-attention tiles from each class were used for cell type quantification (using HoVer-Net)^27^. Attention analysis at the pixel level was performed using Integrated Gradients on the stacked UNI2 and PATH-IC models together^28^. Integrated Gradients works by modifying each pixel in the input image to a baseline, in our case changing the pixel color to white, then quantifying the effect of this change on the prediction of the class. Pixel attention attribution scores were normalized across all class-distinctive high-attention tiles. Finally, attention analysis at the WSI level was performed by computing every tile’s instance embedding PC1 value and overlaying these values on the histology WSI. For clustering analysis, tile instance embedding PC1 values were binned to five groups that represented the distribution of PC1 values that we observed in high-attention score tiles.

### Perturbational Analysis

Perturbational model interpretation of PATH-IC was performed by first selecting a starting tile near the extreme of the instance embedding PC1 axis. We then applied a gradient-descent based approach that in a step-wise fashion perturbed the UNI2 embedding of the start tile and encouraged the perturbed embedding to have an instance embedding PC1 value slightly closer to the opposite extreme. This was repeated until the perturbed embedding passed a predefined PC1 threshold on the other extreme of the instance embedding PC1 axis. Next, intermediate locations in instance embedding space were identified by interpolating between the initial start tile and the perturbed end tile, then converted to UNI2 embeddings via a linear regression model. Finally, the start embedding, perturbed end embedding, and each intermediate embedding were visualized using CRAWFORD to define a path from one high-attention extreme to the other.

### Attention-Based Single Cell Spatial Transcriptomics Analysis

To identify gene expression differences between the cancer cells of High Risk and Typical Risk tumors, we used an internal single cell spatial transcriptomics (scST) dataset. This dataset consists of core biopsies from 204 ER+/HER2-patient samples from the H12O cohort which were selected by an expert breast pathologist, embedded into Formalin Fixed Paraffin Embedded (FFPE) blocks and arrayed onto 9 tissue microarrays (TMAs) with two cores from each patient. The TMA sections for scST profiling were taken roughly 25μm from the sections imaged for histology. The H12O cohort is originally from a study of untreated primary breast cancer patients initiated 25 years ago, and patient follow-up data was recorded including relapse status^34^. These TMAs were profiled with the CosMx SMI RNA Human 6K Discovery Panel and applied to our in-house analysis pipeline. Briefly, after cell segmentation using CellPose^35^ with adapted hyperparameters for human tissue, we applied cell level filtering to select only the cells with more than 100 transcripts per cell. Then, gene expression matrices were normalized by total transcript counts per cell through Seurat V3 using the method SCTransform V2^36^. We used Seurat’s anchors approach to integrate the 9 different TMAs together and remove batch effects. Finally, we used the label transfer approach singleR (v2.0.0)^37^ for cell type classification. To perform attention-based scST analysis, we deployed PATH-IC on the histology images of the TMA, identified the tiles with the most extreme instance embedding PC1 values and stored the tile coordinates. We subset the scST data by cells whose centroid was within the class-distinctive high-attention tile regions on the coregistered coordinate axis and whose cell type was annotated as cancer. We formed pseudo-bulk expression profiles by aggregating the raw counts of cancer epithelial cells from each tile using the *AggregateExpression()* of the Seurat (v4.3.0) R package. We used the limma (v3.54.2) and edgeR (v3.40.2) R packages for normalization and differential expression analyses. Briefly, we used TMM and voom to handle the library size and *voomWithQualityWeights()* function to down-weight low cellularity samples in the normalization. We performed batch correction by assigning the TMA as the major source of batch effect in the gene expression data, as well as cellularity and detection rate as covariates in the *removeBatchEffect()* function. The differential analysis was performed using *lmFit()* and *eBayes()* functions across High Risk versus Typical Risk tiles. We performed GSEA using the GO Biological Process 2021 gene set list and the *fgsea* (v1.24.0) R package.

## Data/Code Availability

All TCGA and GTEx histology and genomic sequencing data used are publicly available. Code for BERGERON and CRAWFORD is available on GitHub (https://github.com/cancersysbio/BERGERON and https://github.com/cancersysbio/CRAWFORD), with weights available on HuggingFace (https://huggingface.co/cancersysbio/PATH-IC and https://huggingface.co/cancersysbio/CRAWFORD_UNI2).

