## Supplemental Figures for "Generative AI Enables Breast Cancer Genomic Subtype Prediction from Histology Images"

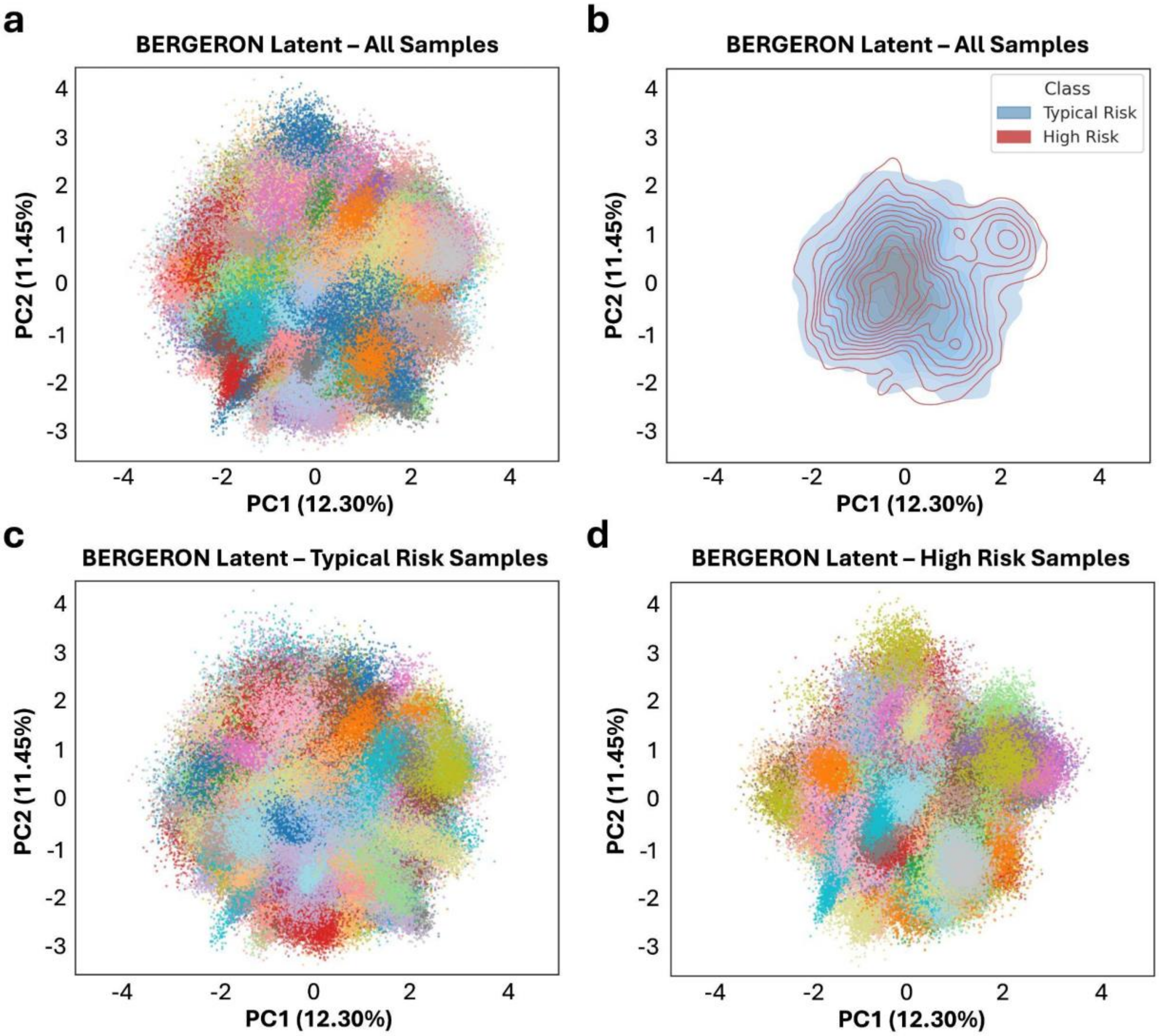


**Supplemental Figure 1. The BERGERON Latent Space.** The latent space of BERGERON, which is defined by the 64-dimensional output of the encoder, clusters tiles from the same WSI together. PCA plots of the UNI2 embeddings of a subset of **a)** all real tiles colored by sample, **b)** all real tiles colored by IC subtype, **c)** all real Typical Risk tiles colored by sample, and **d)** all real High Risk tiles colored by sample. The presence of the class token in the encoder and decoder of BERGERON decouples the latent embeddings, leading to distinct distributions in the latent space for each class and allowing the model to capture better class-specific feature representations.

a

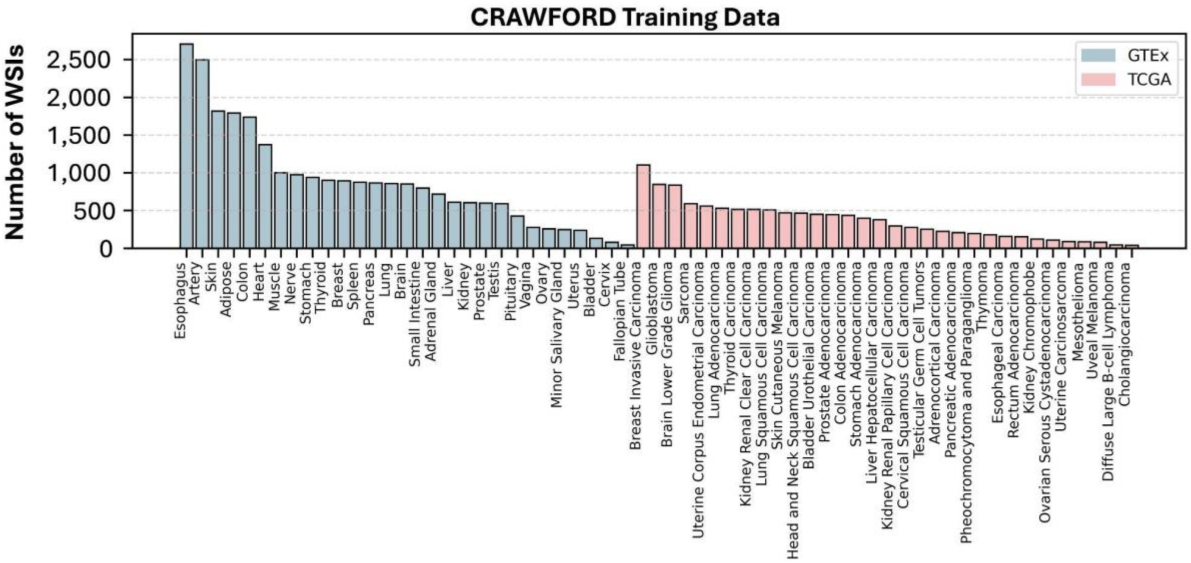


**Supplemental Figure 2. CRAWFORD Training Data. a)** The number of WSIs used from each of the 61 tissue types included in the CRAWFORD training data from the GTEx and TCGA datasets. 100 randomly selected tiles from each WSI were used.

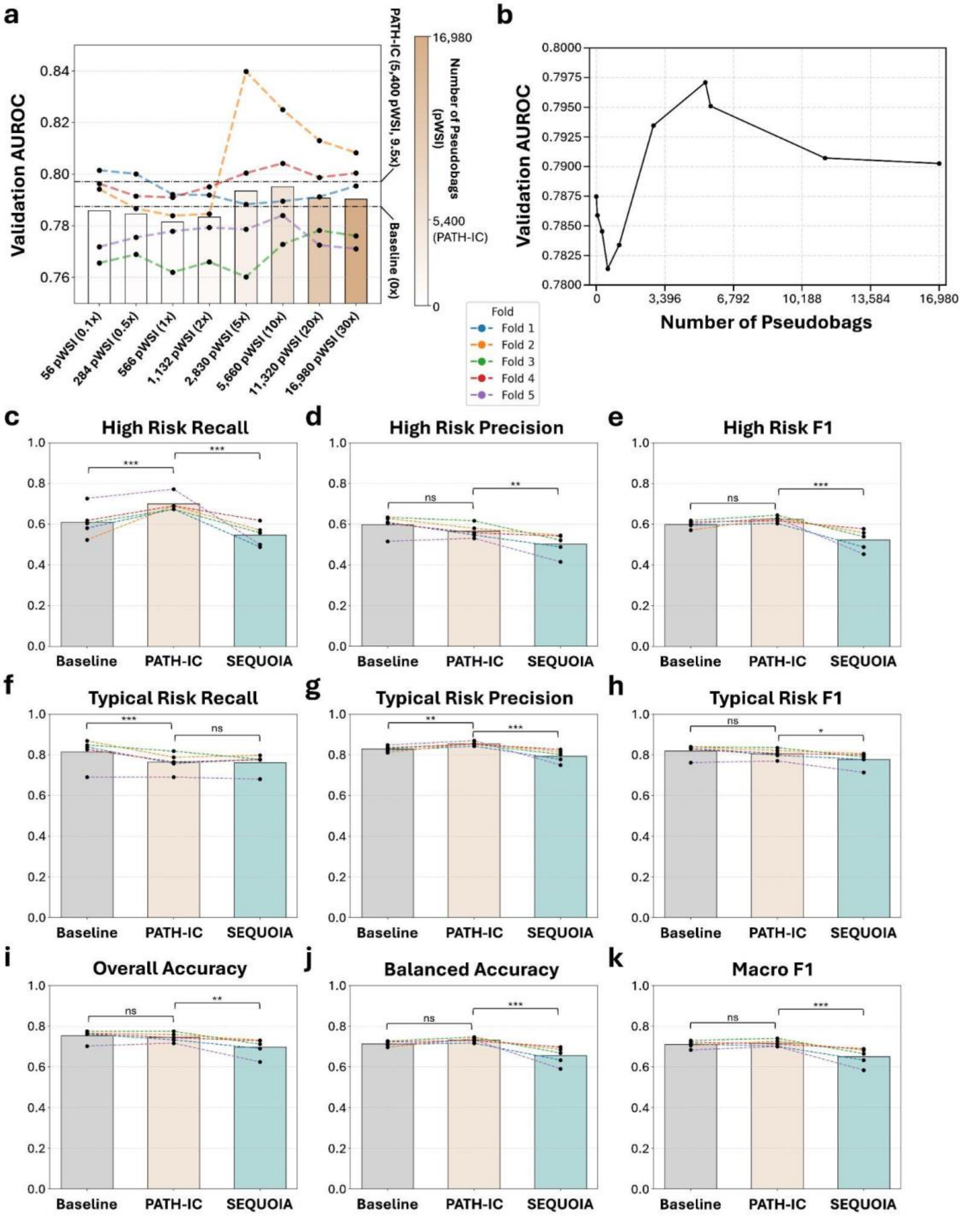


**Supplemental Figure 3. Pseudobag Quantity Tuning and Extended Model Performance. a)** PATH-IC validation AUROC with various quantities of synthetic data. The scalar in parentheses refers to the number of pseudobags used compared to real bags. Matched lines indicate common training and validation folds. **b)** Scatterplot showing PATH-IC mean validation AUROC with increasing numbers of pseudobags. **c-k)** Comparisons of baseline, PATH-IC (trained using the real training data and 5,400 BERGERON-produced pseudobags), and SEQUOIA for predicting IC subtype measured in **c)** High Risk recall, **d)** High Risk precision, **e)** High Risk F1 score, **f)** Typical Risk recall, **g)** Typical Risk precision, **h)** Typical Risk F1 score, **i)** overall accuracy, **j)** balanced accuracy, **k)** and macro F1 score. Matched lines indicate common training and validation folds.

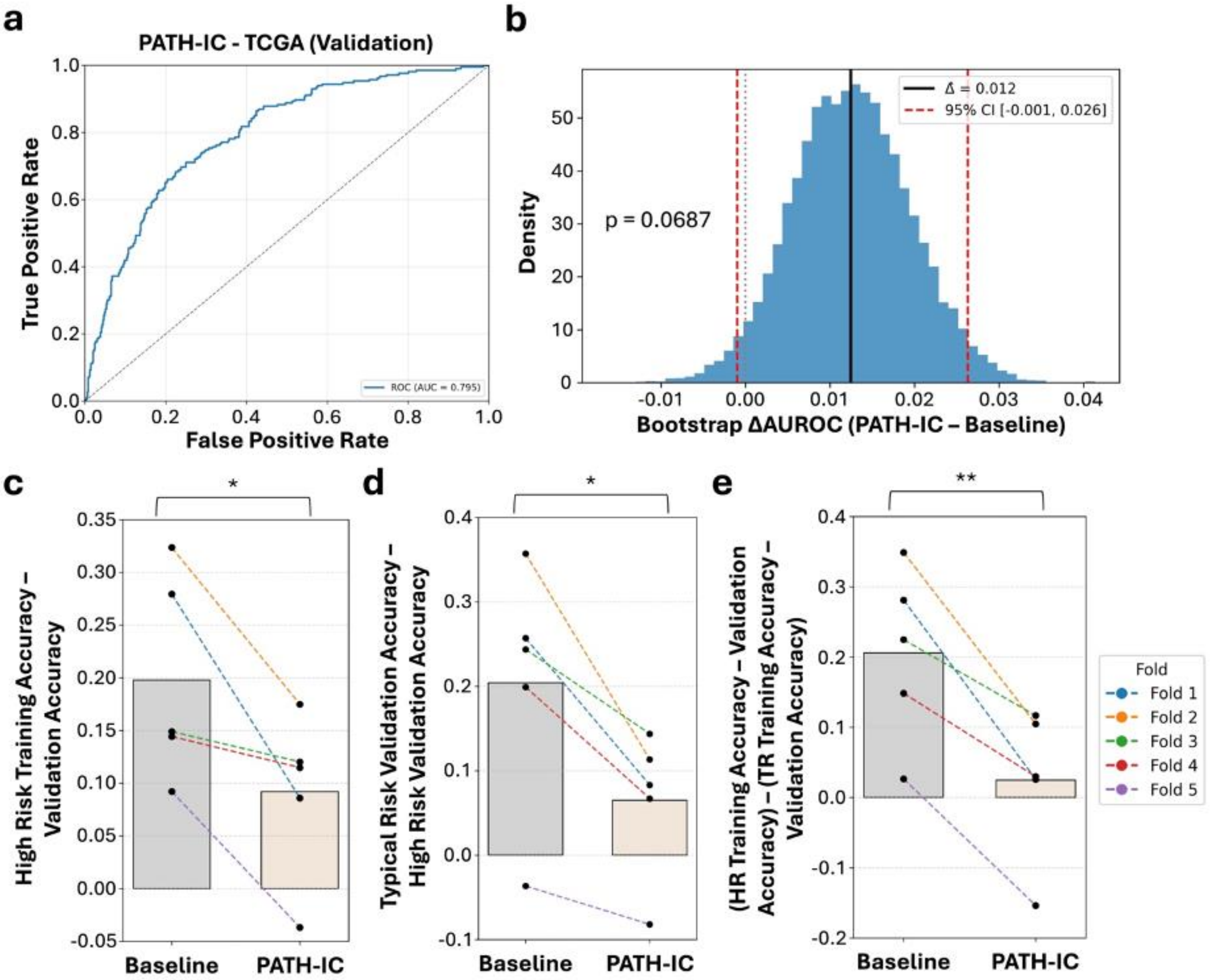


**Supplemental Figure 4. PATH-IC Extended Performance. a)** PATH-IC model validation ROC curve. **b)** Validation sample AUROC bootstrap to measure the change in AUROC between the PATH-IC and the baseline model. **c)** The difference between High Risk Training Accuracy and High Risk Validation Accuracy for Baseline and PATH-IC. **d)** The difference between Typical Risk Validation Accuracy and High Risk Validation Accuracy for Baseline and PATH-IC. **e)** The difference between High Risk Training Accuracy minus High Risk Validation Accuracy and Typical Risk Training Accuracy minus Typical Risk Validation Accuracy for Baseline and PATH-IC.

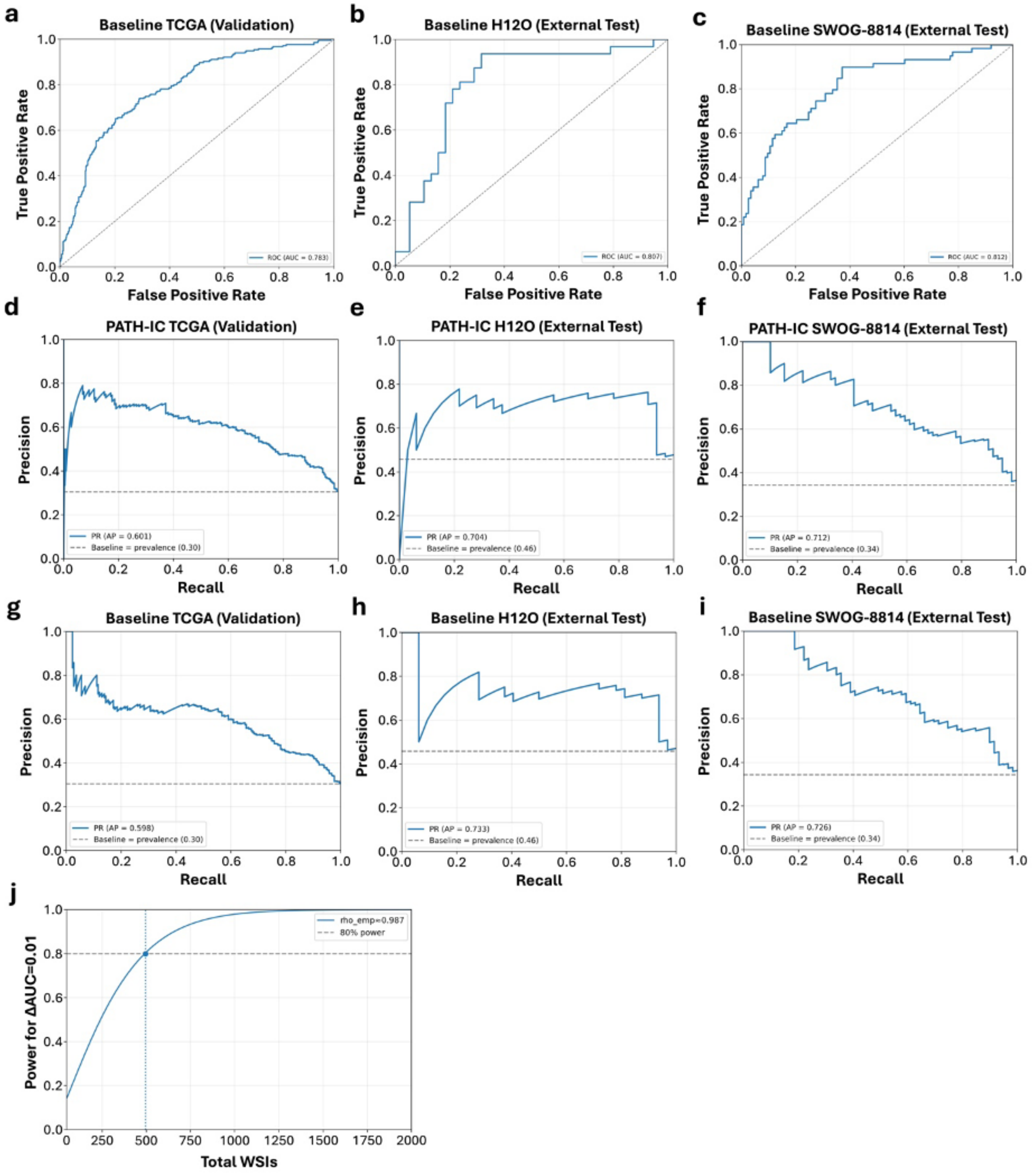


**Supplemental Figure 5. PATH-IC External Validation. a)** Baseline model validation ROC curve. **b)** Baseline model external testing ROC curve for the H12O cohort. **c)** Baseline model external testing ROC curve for the SWOG-8814 cohort. **d)** PATH-IC model validation PRC curve. **e)** PATH-IC model external testing PRC curve for the H12O cohort. **f)** PATH-IC model external testing PRC curve for the SWOG-8814 cohort. **g)** Baseline model validation PRC curve. **h)** Baseline model external testing PRC curve for the H12O cohort. **i)** Baseline model external testing PRC curve for the SWOG-8814 cohort. **j)** Power curve using measured correlation coefficient and a targeted change in AUROC of 0.01.

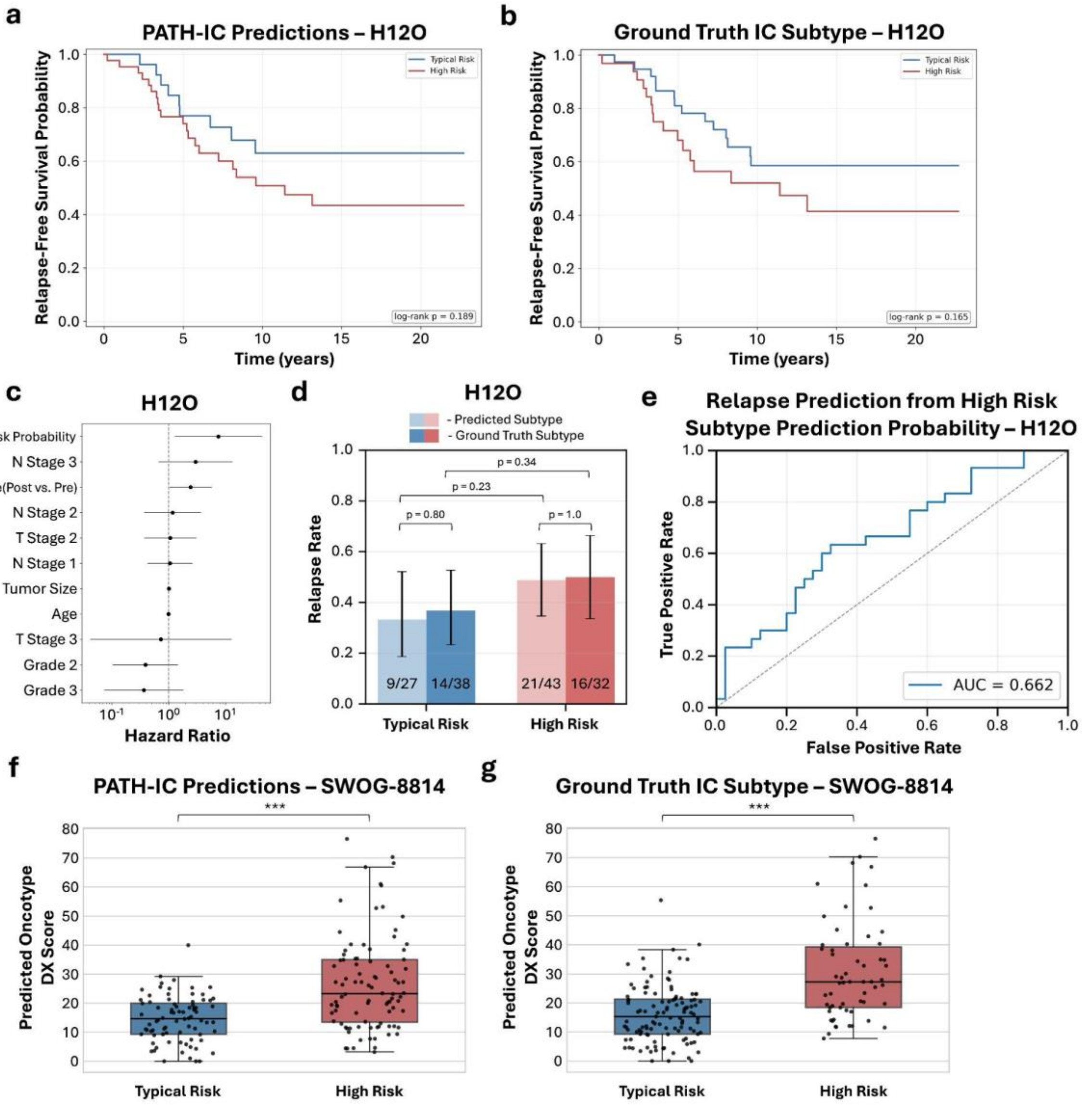


**Supplemental Figure 6. PATH-IC Predictions and Long-Term Relapse. a)** Kaplan-Meier curve for relapse-free survival in the H12O cohort by PATH-IC predicted IC subtype. **b)** Kaplan-Meier curve for relapse-free survival in the H12O cohort by ground truth IC subtype. **c)** Multivariate Cox proportional hazards model measuring the hazard ratio for PATH-IC High Risk prediction probability adjusted for age, menopause status, tumor size, grade, T stage, and N stage in the H12O cohort. **d)** Relapse rate of patients whose samples were predicted to be Typical Risk or High Risk by PATH-IC shown next to relapse rates by ground truth IC subtype in the H12O cohort. **e)** ROC curve for relapse prediction using PATH-IC High Risk prediction probabilities in the H12O cohort. **f)** Inferred Oncotype DX scores separated by PATH-IC predicted IC subtype in the SWOG-8814 cohort. **g)** Inferred Oncotype DX scores separated by ground truth IC subtype in the SWOG-8814 cohort.

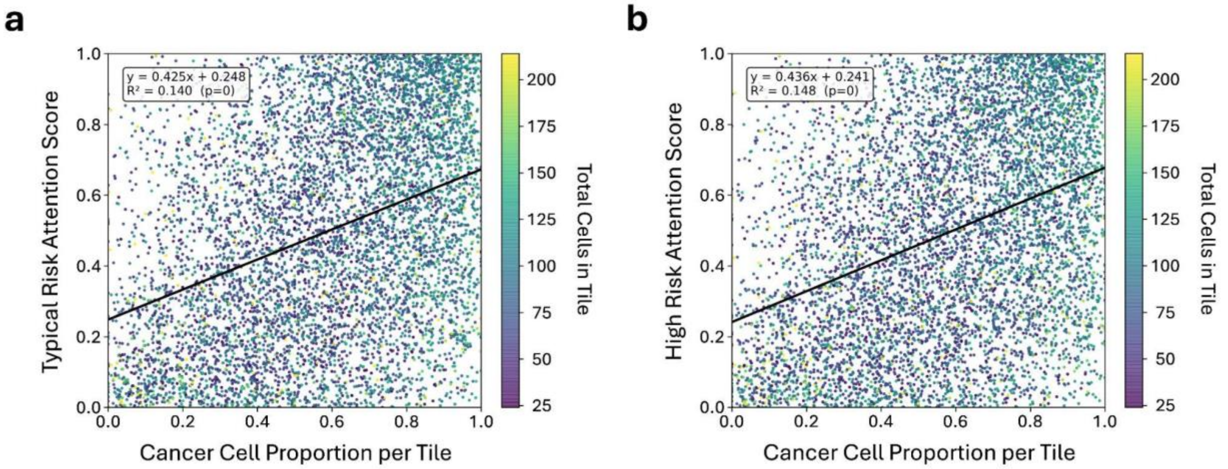


**Supplemental Figure 7. Tile Attention Score Correlates with Cancer Density. a-b)** Scatterplot of the cancer cell proportion per tile and **a)** Typical Risk attention score or **b)** High Risk attention score for 10 randomly selected tiles from each sample. Tiles colored by the total number of cells per tile. Cell counts determined using HoVer-Net.

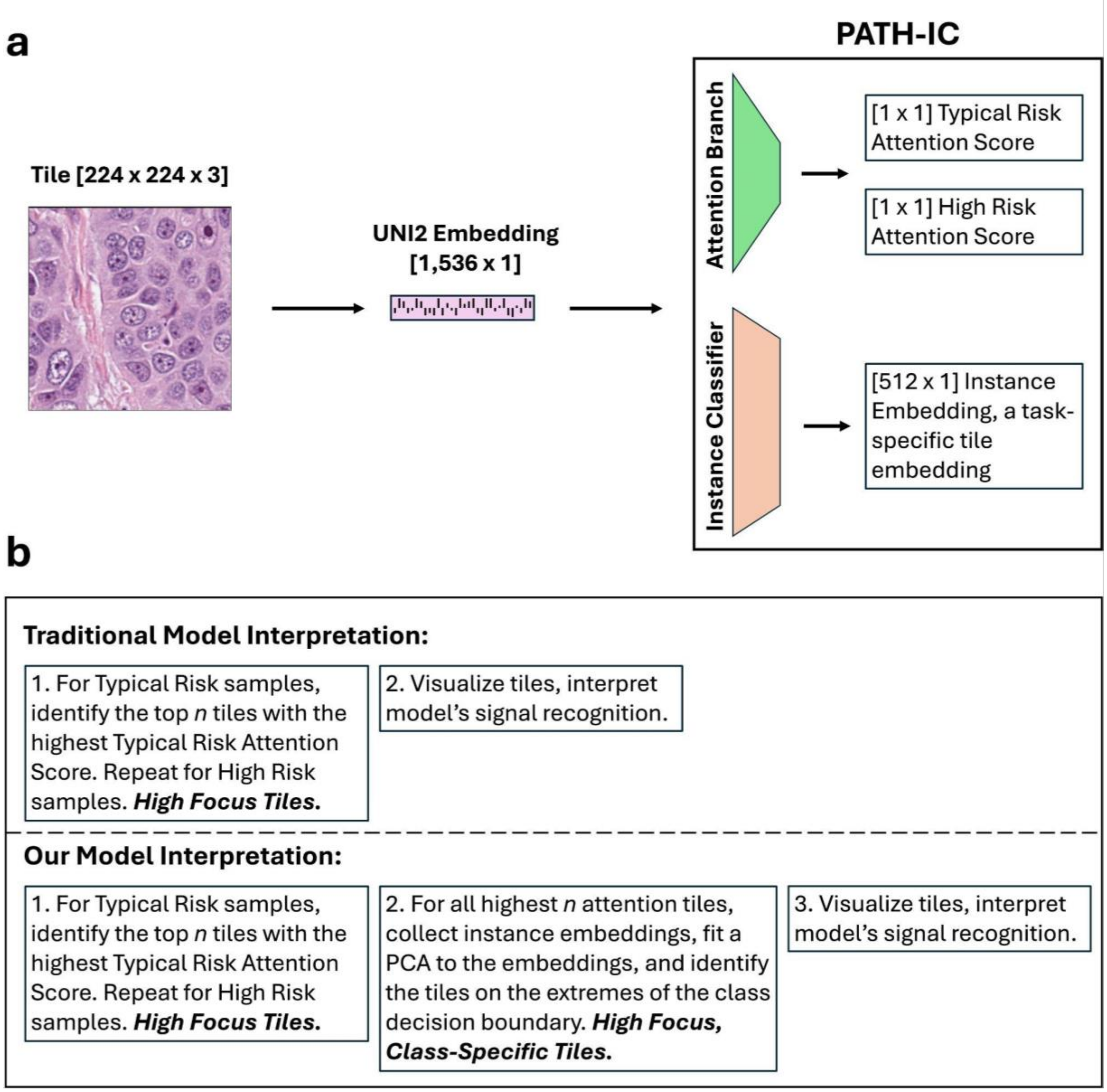


**Supplemental Figure 8. Attention Scores and Instance Embeddings. a)** When a tile is prepared to pass through PATH-IC, it is first converted from an image to a UNI2 embedding. It then passes through the two branches of PATH-IC: the attention branch, which returns a scalar score for each class representing the relative importance of that tile for making a prediction of that class, and the instance classifier branch, which is trained to distinguish tiles from different classes and returns a task-specific tile embedding. **b)** Traditional CLAM model interpretation involves visualizing the tiles in each WSI with the highest attention scores. In addition to conditioning on attention score, we examine the tiles with the most class-specific instance embeddings, as revealed by the PC1 axis of a PCA fit on the instance embeddings of the tiles with the top 100 attention scores per WSI. This reveals the tiles that PATH-IC both focuses most on and finds most class-distinctive.

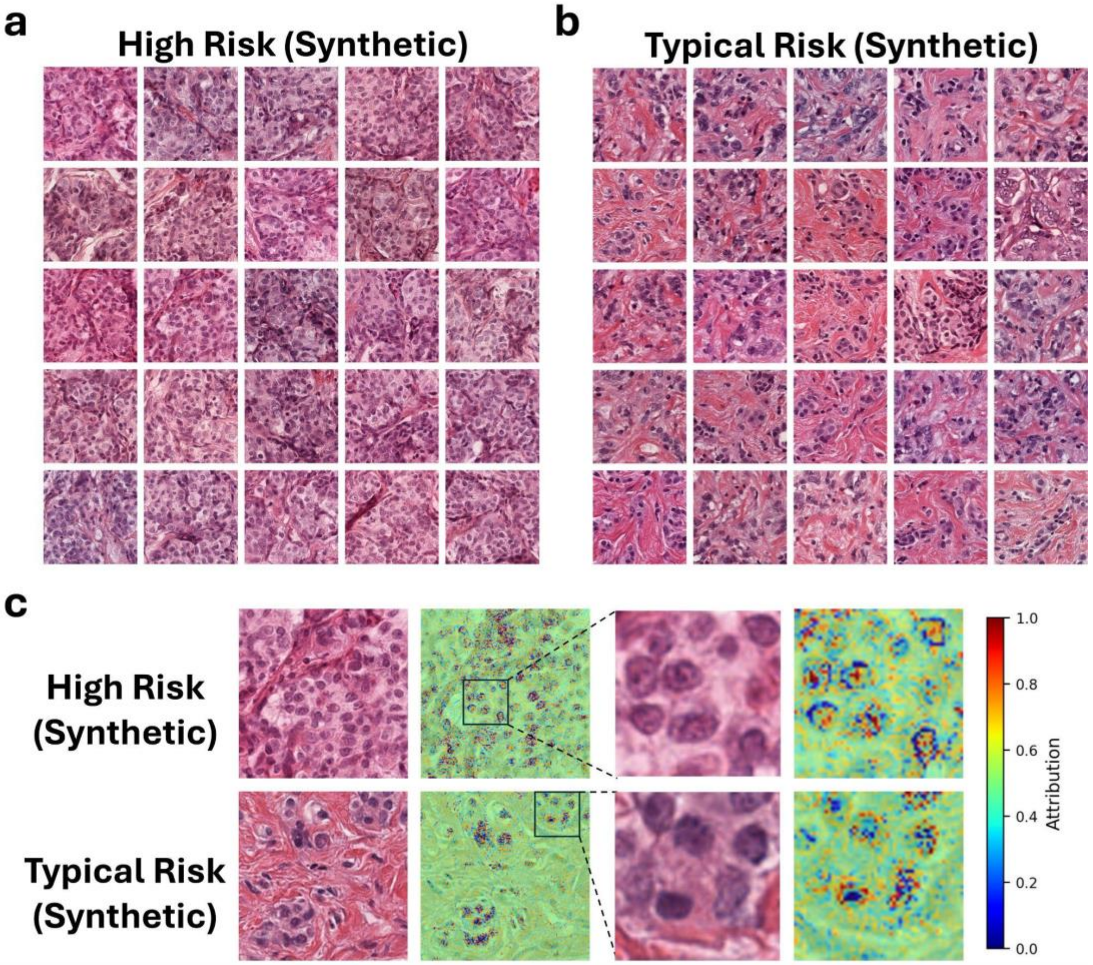


**Supplemental Figure 9. Validation and Interpretation of Synthetic Data. a-b)** Synthetic tiles with the **a)** most negative instance embedding PC1 values (most extreme High Risk) and **b)** most positive instance embedding PC1 values (most extreme Typical Risk) were extracted and visualized using CRAWFORD. **c)** Pixel-level attention assessment using Integrated Gradients.

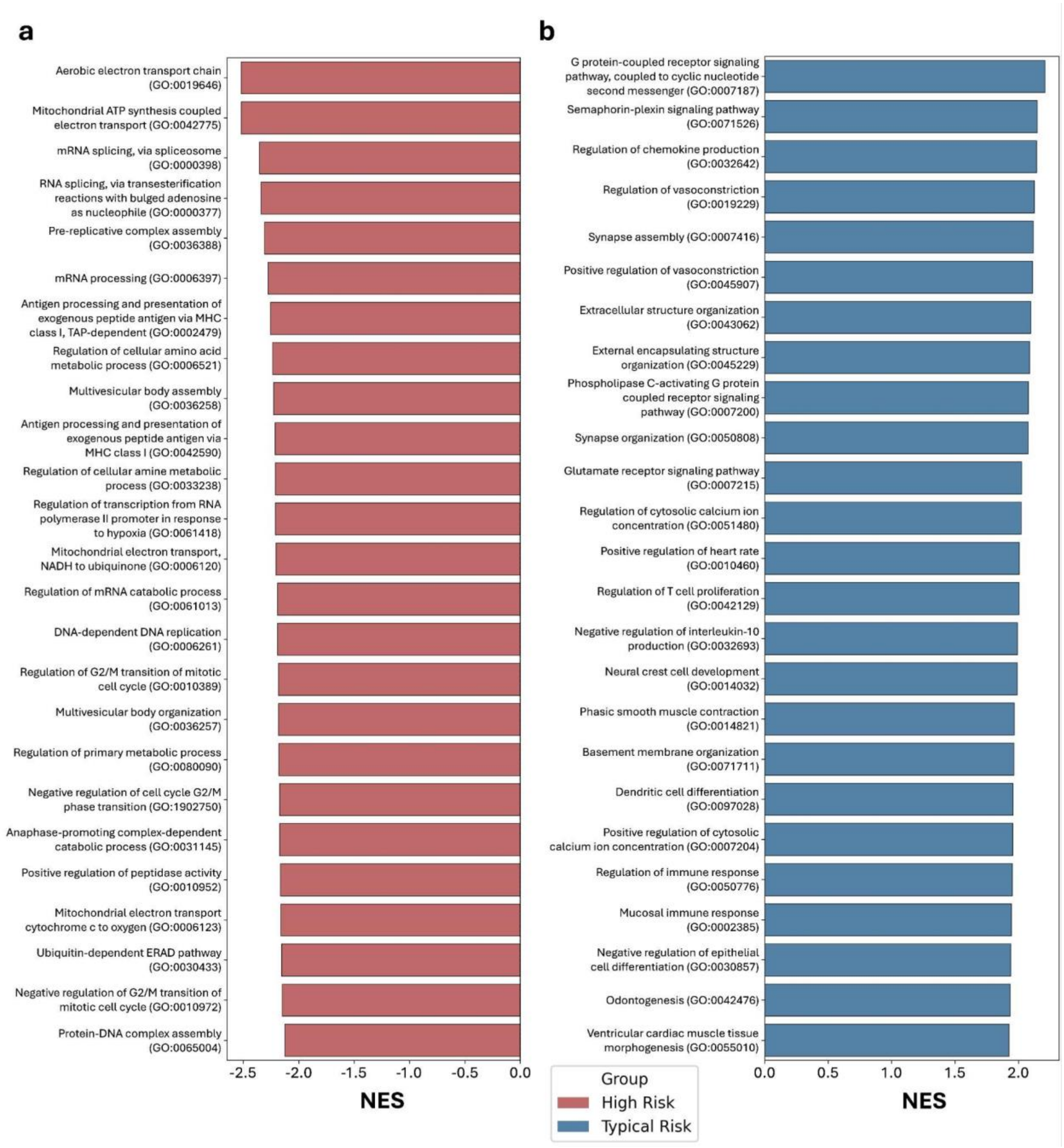


**Supplemental Figure 10. Subtype-Specific Gene Set Enrichment. a-b)** The top 25 enriched gene sets for **a)** High Risk cancer cells and **b)** Typical Risk cancer cells measured by normalized enrichment score (NES).

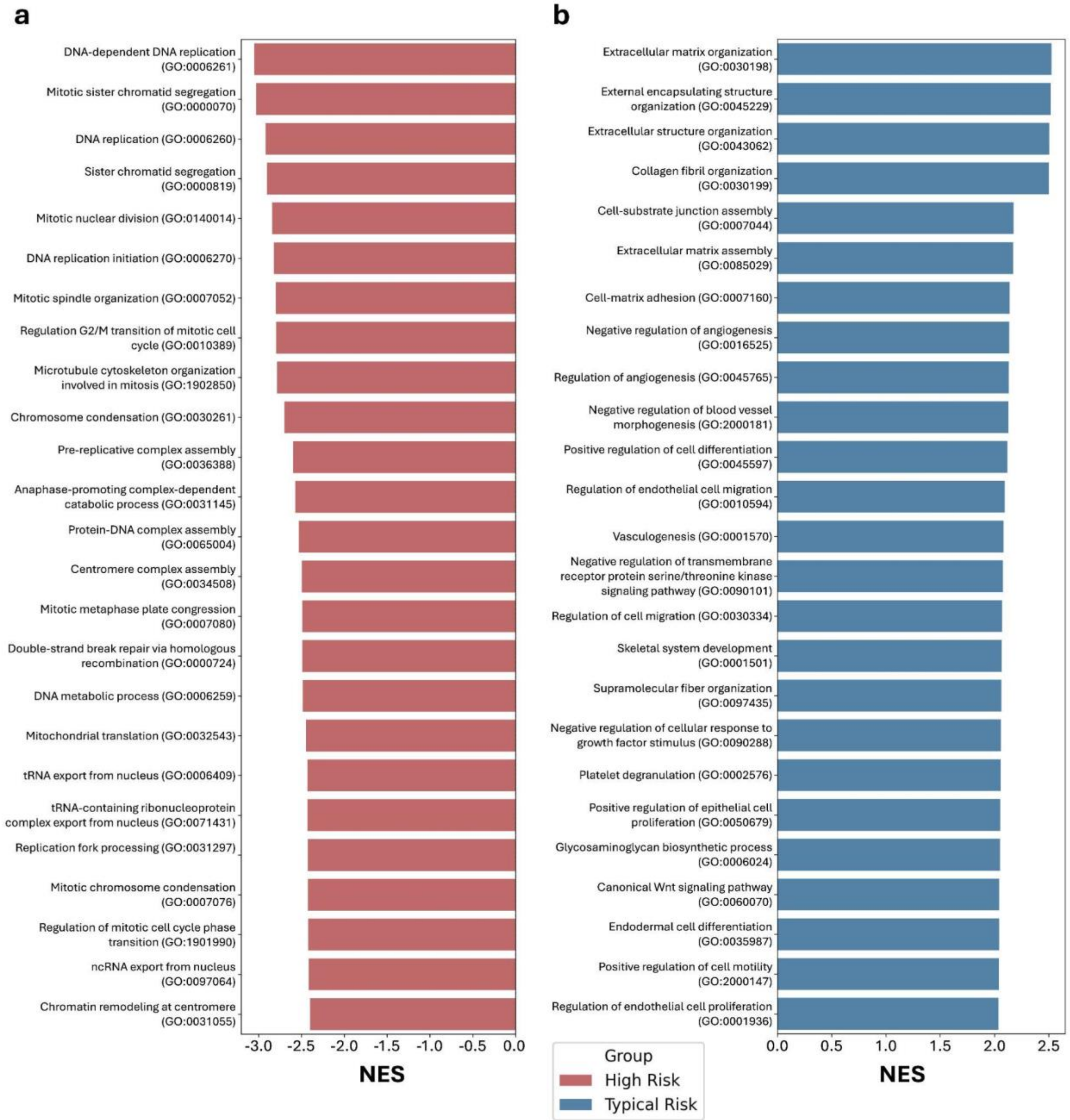


**Supplemental Figure 11. Subtype-Specific Gene Set Enrichment of Imputed RNA expression. a-b)** The top 25 enriched gene sets in imputed bulk mRNA expression from the top 1,000 most class-distinctive high attention tiles for **a)** High Risk cancer cells and **b)** Typical Risk cancer cells measured by normalized enrichment score (NES). Bulk mRNA imputation performed using SEQUOIA on the TCGA-BRCA dataset.

| Parameter | Value |
|---|---|
| Architecture | clam_mb |
| Size | big |
| Dropout | 0.25 |
| Regularization | 1e-5 |
| Learning Rate | 1e-5 |

**Supplemental Table 1. PATH-IC Parameters.** The parameters used to train PATH-IC, a CLAM ABMIL model. All other parameters were default values.

| Parameter | Value |
|---|---|
| Encoder Layers | [512, 256, 128] |
| Decoder Layers | [256, 512, 1024] |
| Dropout | 0.3 |
| Learning Rate | 1e-4 |
| β Initial | 0.01 |
| β Final | 0.05 |
| Latent Dimension Size | 64 |
| Epochs | 50 |

**Supplemental Table 2. BERGERON Parameters.** The parameters used to train BERGERON to create synthetic tile embeddings for PATH-IC training.
